## Supplementary material for "Contrasting cognitive, behavioral, and physiological responses to breathwork vs. naturalistic stimuli in reflective chamber and VR headset environments": S1 Appendix.pdf

### S1 Appendix: Lumena Labs Infinity Cube Spec Sheet

The Lumena Infinity Cube is a sensory-controlled, sensory deprivation, immersive experience tool/instrument/device. The infinity cube is built using custom aluminum extrusions for the structural frame, aerospace-grade mylar mirrored panels for walls and ceiling, a tempered Starphire glass laminated mirror stack for the floor, and a matrix of light-emitting diodes (LEDs) located in all panel seams to create the shape of a cube. Additional LEDs are dispersed in the wall panels as pinpoints of light to give the effect of stars.

#### Usage

The Infinity Cube is designed to provide experiential access to therapeutic states, sensory deprivation, and an enhanced skills learning environment. Utilizing studied and proven modalities such as Breathwork, R.E.S.T., and Chromotherapy.

##### Mechanical Assembly

Infinity Cube is comprised of _ main mechanical components

- Octanorm aluminum extrusion frame
- Aerospace-grade Mylar panels
- Tempered Starphire laminated mirrored floor
- LED 12vDC RGB strip lighting with WS2815 integrated circuit
- Printable outer skins
- Self serve user interface touch screen terminal
- Wired and wireless sound options (headphones, neck speakers)

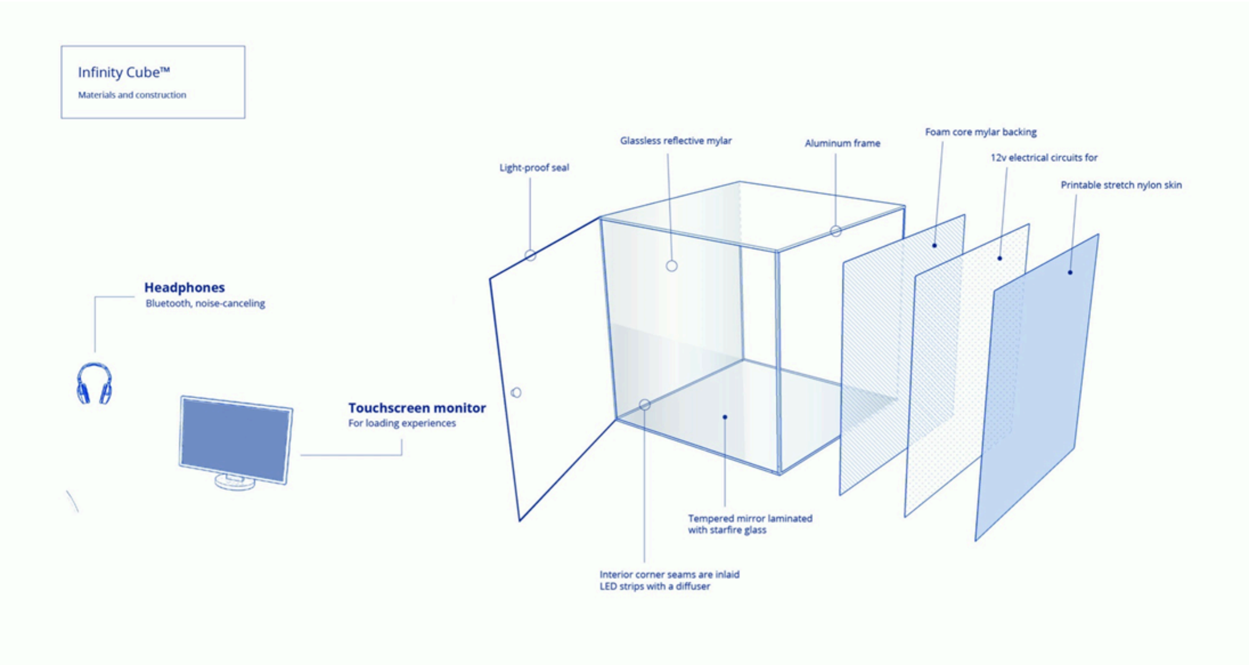


#### Overall System Architecture

##### Information/Data Flow

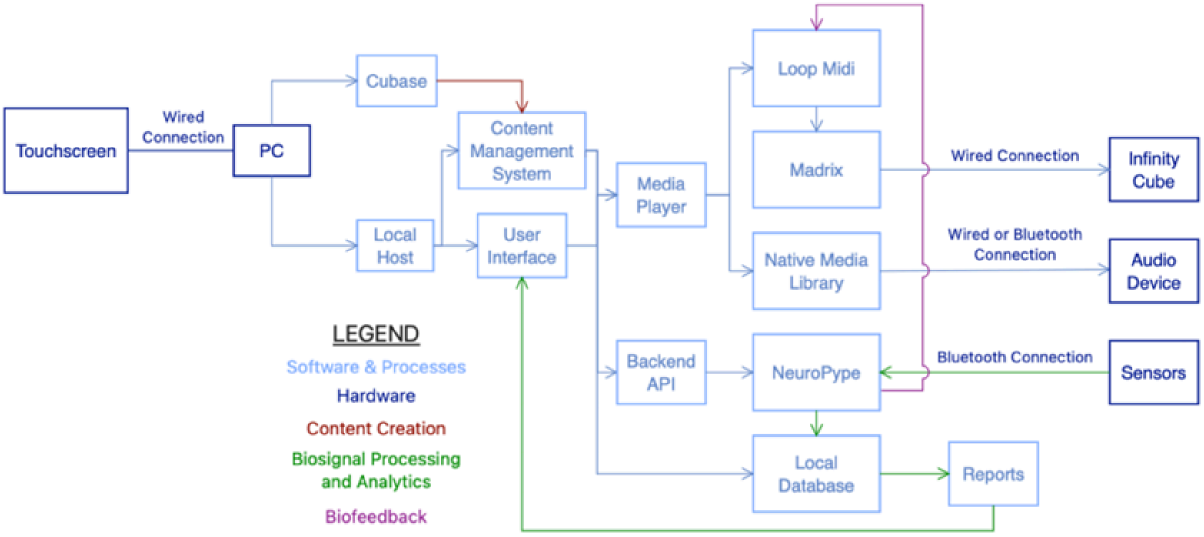


**Composition and Control Software**

All experiences/sessions are composed in Cubase, a digital audio workstation (DAW), where audio is imported and mastered. An accompanying MIDI track of signals/notes is written to match musical rhythm or visual intention of session.

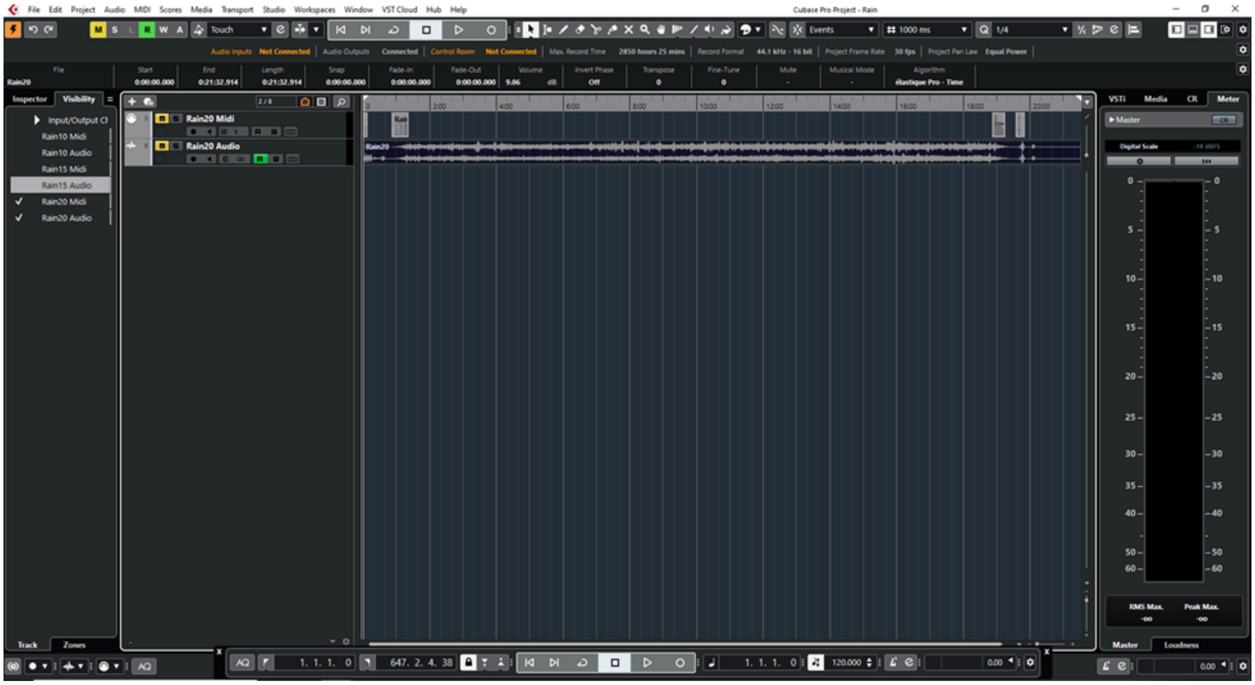

Once a session is finalized in Cubase it is then exported with separate .wav and .midi files. These files are then uploaded and saved into the content management software (CMS).

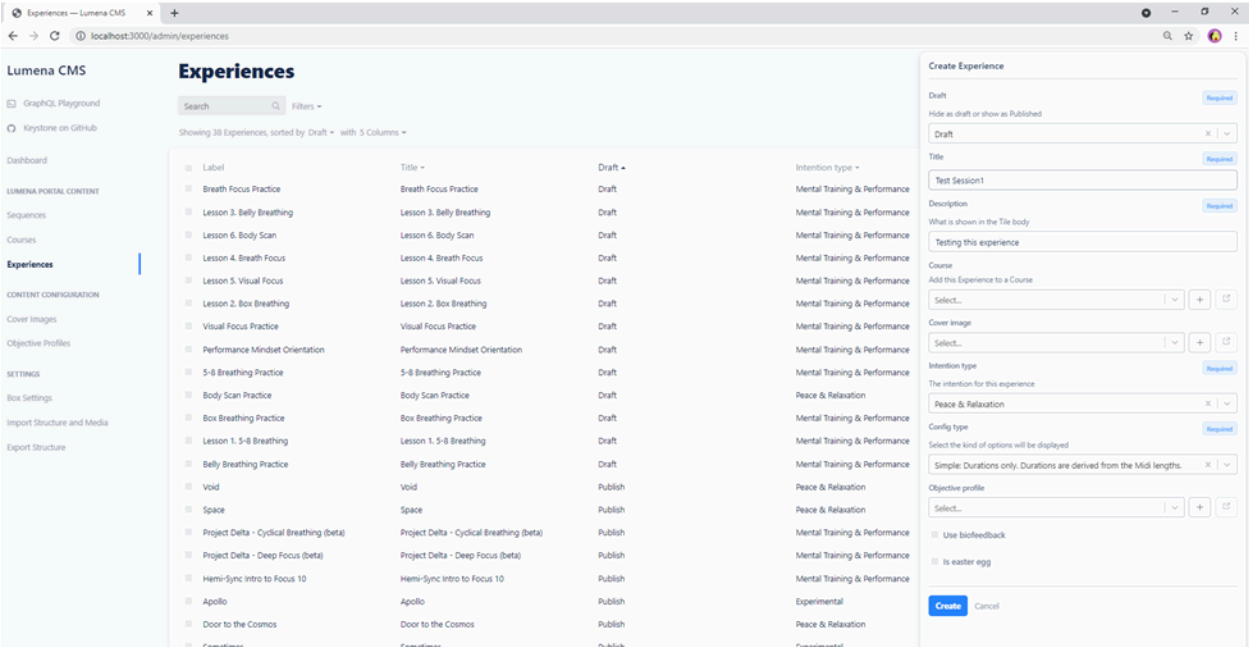


Once session content is uploaded to the CMS sessions are labeled, described, and categorized. Sessions can be made into a single experience or a series using the create course feature. Sessions are now available in the content menu of the user interface and are categorized by intention.

**User Interface**

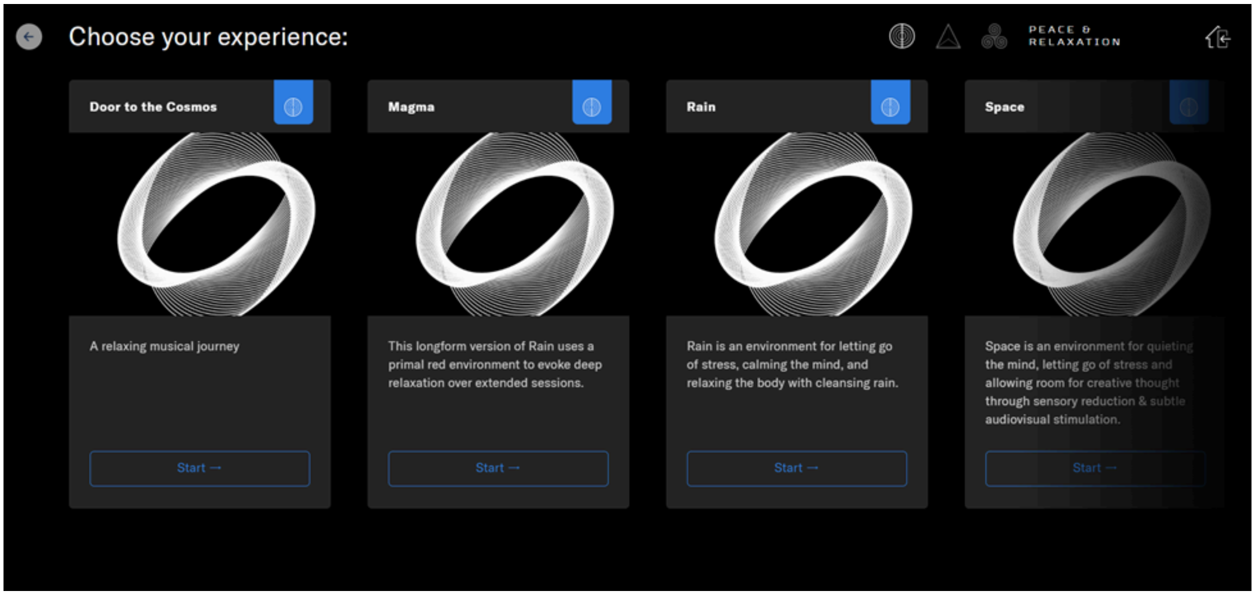


Selecting a session starts a countdown in which the CMS sends the queued MIDI data to loopMIDI and then to Madrix for control of all lighting. The saved audio file is also queued and sent to the local media library player. Files are synchronized and at the end of the start timer, the session begins. The user has a 30-second timer to enter the Cube, sit down in the chair and prepare for the session to start. During this time the Cube is illuminated for safety and orientation of the space.

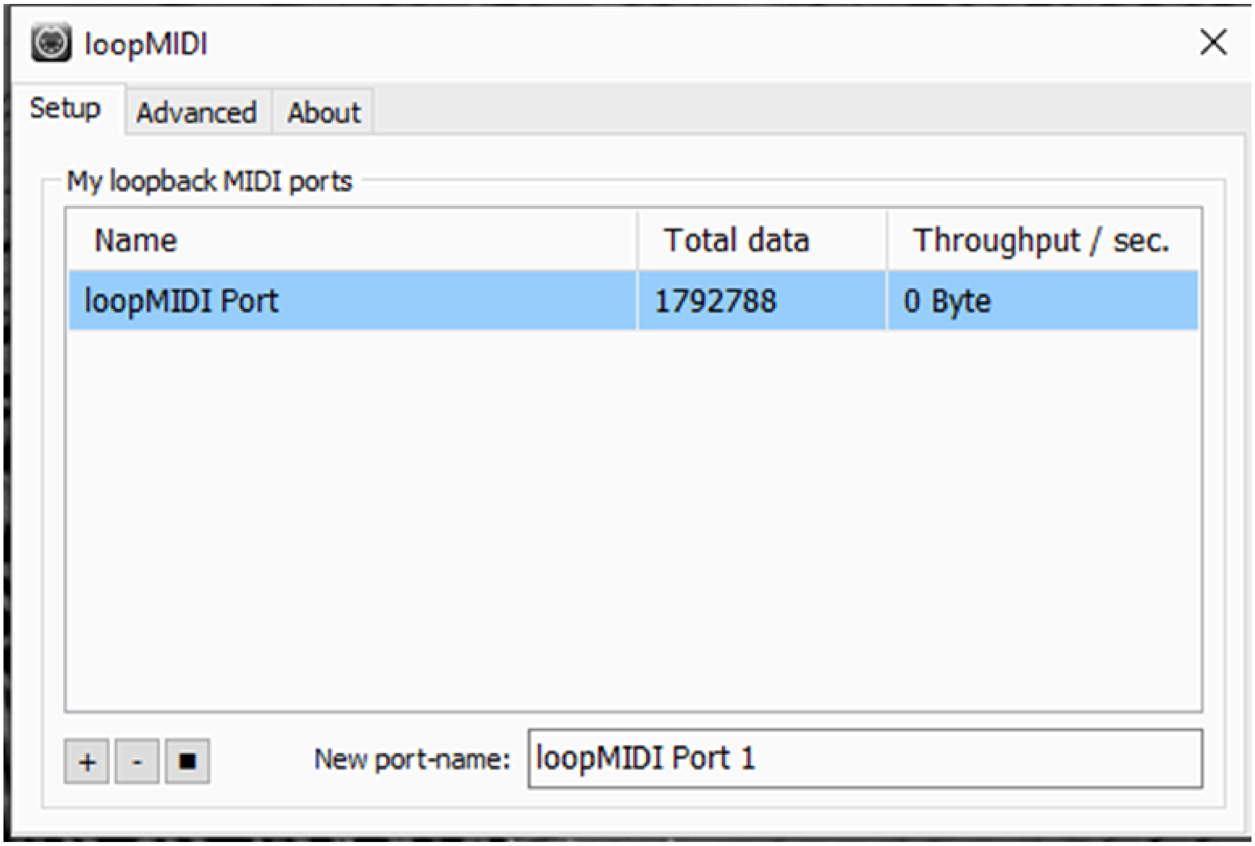


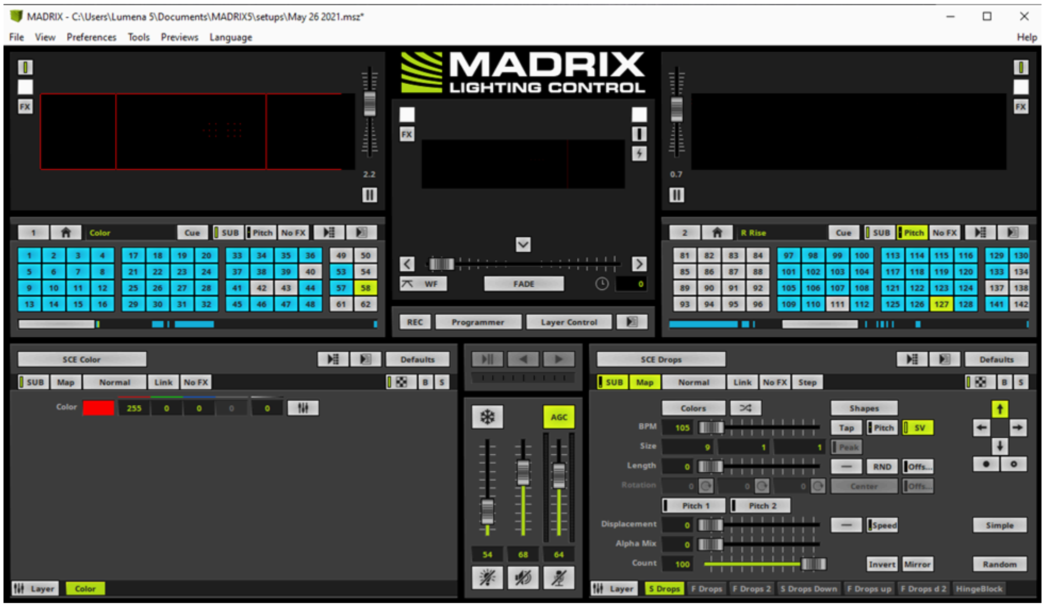


At the completion of a session the user exits the Cube and proceeds back to the terminal to either 1. Without the use of biosensors, end the session and return to the home menu. 2. Using biosensors, wait a moment for the data collected during the session to be processed through our integrated use of Nueropype (biometrics and neurometrics processing suite) to generate a report of biometrics and task metrics for that session. After reviewing the report, the user can end the session and return to the main menu.

#### Infinity Cube Hardware

##### LED Layout and Controls

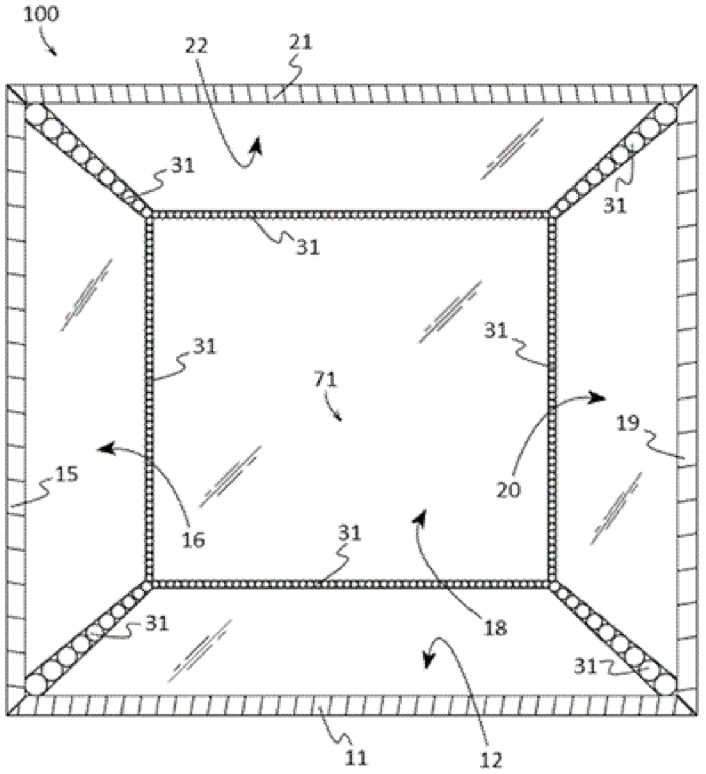


*FIG. 6*

LEDs are mounted inside the frame in all seams to create an illuminated cube.

**Integrated Peripherals**

-Cognionics Quick 20 Dry EEG headset (Q20). The Q20 is an FDA cleared 20 physiological acquisition channel, 1 EEG reference signal acquisition dry headset. This device has a sample rate of 500 per second, a bandwidth of 0-131 Hz with true DC coupling, 10 meter Bluetooth low energy wireless range, and a battery runtime of 6 hours (uses 2 AAA rechargeable batteries). This device is used to collect neurological data from user to report cognitive changes during sessions as well as, on certain sessions, used as a brain computer interface controller to live manipulate the immersive experience inside the cube during a session. LEDs are

-Zephyr BioHarness 3.0 chest strap. The BioHarness is a physiological monitoring telemetry device capable of collecting and streaming data on ECG, heart rate, and RR. This device uses BTLE to stream data, has an internal rechargeable lithium polymer battery (max runtime of 24hrs per charge), ECG sampling frequency of 250 Hz, and breathing rate sampling frequency of 25 Hz. This device is used to collect physiological data from user to report changes in HR, RR, and HRV during sessions as well as, on certain sessions, used as a contoller to live manipulate the immersive experience inside the cube during a session.

**Electrical Specifications**

110v standard electrical outlet is required to power the Infinity Cube.
Two 12v Mean Well power supplies provide constant voltage to LEDs enabling proper dimming control for lighting system in low light sessions.

Two 10amp inline fuse silicon wires run to two power distribution blocks located on bottom left back of cube and top right front of cube. This enables an even distribution of power to LEDs.
Main power consumption is from terminal computer, LED power supplies run at ≤ 50% capacity during most sessions. LED MIDI data is sent from Madrix lighting software to Madrix Nebulas for processing and then sent to LEDs inside the cube.
