## Supplementary material for "Contrasting cognitive, behavioral, and physiological responses to breathwork vs. naturalistic stimuli in reflective chamber and VR headset environments": S1 Tables A-I.pdf

### S1: Timepoint effects within groups

#### Table 1A (MindGym Breathwork)

| Group (MindGym Breathwork) | | | | | | | | | | | | | |
|---|---|---|---|---|---|---|---|---|---|---|---|---|---|
| | Pre | | | Post | | | Wilcoxon Signed-Rank test | | | | | | |
| DV | n | M | SD | n | M | SD | W | Z | p.raw | p.fdr | RBC | RBC (low) | RBC (high) |
| TMT (RTACC) | 30 | 135.19 | 57.76 | 30 | 137.86 | 150.46 | 337.00 | 2.15 | 0.031 | 0.0543 | 0.45 | 0.07 | 0.71 |
| Architex (Total Speed) | 30 | 193.91 | 87.87 | 30 | 142.14 | 60.94 | 396.00 | 3.36 | < .001 | 0.0007 | 0.70 | 0.43 | 0.86 |
| Architex (Total Score) | 30 | 39.07 | 3.22 | 30 | 38.13 | 3.34 | 258.50 | 1.67 | 0.097 | 0.1132 | 0.37 | -0.05 | 0.67 |
| Anxiety (STAI) | 31 | 35.45 | 8.63 | 31 | 31.77 | 8.86 | 323.50 | 2.29 | 0.022 | 0.0513 | 0.49 | 0.12 | 0.74 |
| Arousal | 31 | 2.61 | 1.52 | 31 | 2.42 | 1.46 | 130.50 | 0.52 | 0.609 | 0.6090 | 0.13 | -0.34 | 0.55 |
| Valence | 31 | 6.71 | 1.87 | 31 | 7.74 | 2.03 | 57.00 | -2.03 | 0.042 | 0.0588 | -0.51 | -0.78 | -0.07 |
| POMS | 31 | 12.45 | 24.28 | 31 | 2.55 | 22.53 | 331.00 | 2.92 | 0.004 | 0.0140 | 0.63 | 0.31 | 0.82 |

**Table 1A.** Nonparametric t-test (Wilcoxon Signed-Rank) of timepoint (pre vs post) differences for the MindGym (Breathwork) group. 'DV' refers to dependent variable. 'n' refers to group sample size. 'M' and 'SD' refer to mean and standard deviation. 'p.raw' and 'p.fdr' refer to the uncorrected p value and FDR-corrected p value. 'RBC' refers to rank biserial correlation as a measure of effect size (ranging from -1 to 1) and reflects the relative proportions of positive and negative ranks. 'RBC (low)' and 'RBC (high)' refer to the lower and upper bounds of the 95% confidence interval of RBC.

#### Table 1B (MindGym Rain)

| Group (MindGym Rain) | | | | | | | | | | | | | |
|---|---|---|---|---|---|---|---|---|---|---|---|---|---|
| | Pre | | | Post | | | Wilcoxon Signed-Rank test | | | | | | |
| DV | n | M | SD | n | M | SD | W | Z | p.raw | p.fdr | RBC | RBC (low) | RBC (high) |
| TMT (RTACC) | 31 | 169.98 | 113.92 | 31 | 127.34 | 53.43 | 411.00 | 3.19 | < .001 | 0.0002 | 0.66 | 0.37 | 0.83 |
| Architex (Total Speed) | 30 | 189.86 | 78.98 | 30 | 116.97 | 42.04 | 421.00 | 4.40 | < .001 | 0.0002 | 0.94 | 0.86 | 0.97 |
| Architex (Total Score) | 30 | 37.40 | 7.53 | 30 | 38.05 | 7.93 | 99.50 | -1.44 | 0.152 | 0.1773 | -0.34 | -0.67 | 0.11 |
| Anxiety (STAI) | 32 | 37.59 | 9.01 | 32 | 33.50 | 6.08 | 353.50 | 2.94 | 0.003 | 0.0053 | 0.63 | 0.31 | 0.82 |
| Arousal | 32 | 3.06 | 2.30 | 32 | 2.69 | 1.75 | 140.50 | 0.46 | 0.657 | 0.6570 | 0.11 | -0.35 | 0.53 |
| Valence | 32 | 7.44 | 1.50 | 32 | 7.84 | 1.94 | 75.50 | -1.90 | 0.054 | 0.0756 | -0.45 | -0.74 | -0.02 |
| POMS | 32 | 15.47 | 22.05 | 32 | 4.97 | 16.69 | 442.00 | 3.80 | < .001 | 0.0002 | 0.78 | 0.57 | 0.90 |

**Table 1B.** Nonparametric t-test (Wilcoxon Signed-Rank) of timepoint (pre vs post) differences for the MindGym (Rain) group. 'DV' refers to dependent variable. 'n' refers to group sample size. 'M' and 'SD' refer to mean and standard deviation. 'p.raw' and 'p.fdr' refer to the uncorrected p value and FDR-corrected p value. 'RBC' refers to rank biserial correlation as a measure of effect size (ranging from -1 to 1) and reflects the relative proportions of positive and negative ranks. 'RBC (low)' and 'RBC (high)' refer to the lower and upper bounds of the 95% confidence interval of RBC.

#### Table 1C (VR Breathwork)

| Group (VR Breathwork) | | | | | | | | | | | | | |
|---|---|---|---|---|---|---|---|---|---|---|---|---|---|
| | Pre | | | Post | | | Wilcoxon Signed-Rank test | | | | | | |
| DV | n | M | SD | n | M | SD | W | Z | p.raw | p.fdr | RBC | RBC (low) | RBC (high) |
| TMT (RTACC) | 33 | 195.94 | 200.00 | 31 | 152.73 | 96.38 | 329.00 | 1.59 | 0.116 | 0.2030 | 0.33 | -0.06 | 0.63 |

| | | | | | | | | | | | | |
|---|---|---|---|---|---|---|---|---|---|---|---|---|
| Architex (Total Speed) | 27 | 246.99 | 300.40 | 27 | 132.03 | 37.98 | 355.00 | 3.99 | < .001 | 0.0004 | 0.88 | 0.73 | 0.95 |
| Architex (Total Score) | 27 | 38.30 | 3.31 | 27 | 37.93 | 3.65 | 188.00 | 1.09 | 0.282 | 0.3290 | 0.25 | -0.20 | 0.62 |
| Anxiety (STAI) | 33 | 36.64 | 11.01 | 33 | 30.94 | 7.31 | 405.00 | 3.55 | < .001 | 0.0004 | 0.74 | 0.50 | 0.88 |
| Arousal | 33 | 2.55 | 1.82 | 33 | 3.24 | 2.11 | 135.00 | -1.30 | 0.195 | 0.2730 | -0.29 | -0.62 | 0.14 |
| Valence | 33 | 7.46 | 1.39 | 33 | 7.33 | 2.80 | 176.50 | -0.30 | 0.772 | 0.7720 | -0.07 | -0.46 | 0.35 |
| POMS | 33 | 12.79 | 29.37 | 33 | 2.21 | 23.77 | 406.00 | 3.10 | 0.002 | 0.0047 | 0.64 | 0.34 | 0.82 |

**Table 1C.** Nonparametric t-test (Wilcoxon Signed-Rank) of timepoint (pre vs post) differences for the VR (Breathwork) group. 'DV' refers to dependent variable. 'n' refers to group sample size. 'M' and 'SD' refer to mean and standard deviation. 'p.raw' and 'p.fdr' refer to the uncorrected p value and FDR-corrected p value. 'RBC' refers to rank biserial correlation as a measure of effect size (ranging from -1 to 1) and reflects the relative proportions of positive and negative ranks. 'RBC (low)' and 'RBC (high)' refer to the lower and upper bounds of the 95% confidence interval of RBC.

#### Table 1D (VR Rain)

| Group (VR Rain) | | | | | | | | | | | | | |
|---|---|---|---|---|---|---|---|---|---|---|---|---|---|
| | Pre | | | Post | | | Wilcoxon Signed-Rank test | | | | | | |
| **DV** | **n** | **M** | **SD** | **n** | **M** | **SD** | **W** | **Z** | **p.raw** | **p.fdr** | **RBC** | **RBC (low)** | **RBC (high)** |
| TMT (RTACC) | 30 | 161.35 | 203.67 | 30 | 125.93 | 91.44 | 308.00 | 1.55 | 0.124 | 0.3523 | 0.33 | -0.07 | 0.63 |
| Architex (Total Speed) | 28 | 217.82 | 89.32 | 28 | 148.57 | 67.71 | 379.00 | 4.01 | < .001 | 0.0007 | 0.87 | 0.72 | 0.94 |
| Architex (Total Score) | 28 | 38.34 | 2.58 | 28 | 37.38 | 3.47 | 197.50 | 0.94 | 0.352 | 0.4928 | 0.22 | -0.23 | 0.58 |
| Anxiety (STAI) | 30 | 34.83 | 9.88 | 30 | 33.40 | 6.77 | 235.00 | 0.73 | 0.473 | 0.5518 | 0.16 | -0.26 | 0.53 |
| Arousal | 30 | 3.50 | 2.66 | 30 | 2.87 | 2.16 | 208.50 | 1.24 | 0.215 | 0.3763 | 0.28 | -0.16 | 0.63 |
| Valence | 30 | 7.60 | 1.83 | 30 | 7.63 | 1.73 | 122.50 | -0.13 | 0.908 | 0.9080 | -0.03 | -0.47 | 0.42 |
| POMS | 30 | 13.17 | 31.32 | 30 | 6.00 | 23.07 | 266.50 | 1.45 | 0.151 | 0.3523 | 0.31 | -0.10 | 0.63 |

**Table 1D.** Nonparametric t-test (Wilcoxon Signed-Rank) of timepoint (pre vs post) differences for the VR (Rain) group. 'DV' refers to dependent variable. 'n' refers to group sample size. 'M' and 'SD' refer to mean and standard deviation. 'p.raw' and 'p.fdr' refer to the uncorrected p value and FDR-corrected p value. 'RBC' refers to rank biserial correlation as a measure of effect size (ranging from -1 to 1) and reflects the relative proportions of positive and negative ranks. 'RBC (low)' and 'RBC (high)' refer to the lower and upper bounds of the 95% confidence interval of RBC.

#### Table 1E (MindGym)

| Group (MindGym) | | | | | | | | | | | | | |
|---|---|---|---|---|---|---|---|---|---|---|---|---|---|
| | Pre | | | Post | | | Wilcoxon Signed-Rank test | | | | | | |
| **DV** | **n** | **M** | **SD** | **n** | **M** | **SD** | **W** | **Z** | **p.raw** | **p.fdr** | **RBC** | **RBC (low)** | **RBC (high)** |
| TMT (RTACC) | 61 | 152.87 | 91.70 | 61 | 132.51 | 111.34 | 1470.00 | 3.77 | < .001 | 0.0002 | 0.56 | 0.32 | 0.72 |
| Architex (Total Speed) | 60 | 191.88 | 82.86 | 60 | 129.55 | 53.43 | 1613.00 | 5.50 | < .001 | 0.0002 | 0.82 | 0.70 | 0.90 |
| Architex (Total Score) | 60 | 38.23 | 5.80 | 60 | 38.09 | 6.03 | 702.50 | 0.37 | 0.714 | 0.7140 | 0.06 | -0.25 | 0.36 |
| Anxiety (STAI) | 63 | 36.54 | 8.82 | 63 | 32.65 | 7.57 | 1328.00 | 3.66 | < .001 | 0.0002 | 0.55 | 0.32 | 0.73 |
| Arousal | 63 | 2.84 | 1.95 | 63 | 2.56 | 1.60 | 529.50 | 0.68 | 0.493 | 0.5752 | 0.12 | -0.22 | 0.43 |
| Valence | 63 | 7.08 | 1.72 | 63 | 7.79 | 1.97 | 260.00 | -2.74 | 0.006 | 0.0084 | -0.48 | -0.69 | -0.18 |
| POMS | 63 | 13.98 | 23.03 | 63 | 3.78 | 19.66 | 1516.50 | 4.77 | < .001 | 0.0002 | 0.71 | 0.54 | 0.83 |

**Table 1E.** Nonparametric t-test (Wilcoxon Signed-Rank) of timepoint (pre vs post) differences for the MindGym group. 'DV' refers to dependent variable. 'n' refers to group sample size. 'M' and 'SD' refer to mean and standard deviation. 'p.raw' and 'p.fdr' refer to the uncorrected p value and FDR-corrected p value. 'RBC' refers to rank biserial correlation as a measure of effect size (ranging from -1 to 1) and reflects the relative proportions of positive and negative ranks. 'RBC (low)' and 'RBC (high)' refer to the lower and upper bounds of the 95% confidence interval of RBC.

#### Table 1F (VR)

| Group (VR) | | | | | | | | | | | | | |
|---|---|---|---|---|---|---|---|---|---|---|---|---|---|
| | Pre | | | Post | | | Wilcoxon Signed-Rank test | | | | | | |
| **DV** | **n** | **M** | **SD** | **n** | **M** | **SD** | **W** | **Z** | **p.raw** | **p.fdr** | **RBC** | **RBC (low)** | **RBC (high)** |
| TMT (RTACC) | 63 | 179.47 | 200.88 | 61 | 139.55 | 94.17 | 1253.00 | 2.21 | 0.027 | 0.0473 | 0.33 | 0.05 | 0.56 |
| Architex (Total Speed) | 55 | 232.14 | 218.30 | 55 | 140.45 | 55.28 | 1439.00 | 5.61 | < .001 | 0.0004 | 0.87 | 0.77 | 0.93 |
| Architex (Total Score) | 55 | 38.32 | 2.93 | 55 | 37.65 | 3.54 | 754.50 | 1.41 | 0.158 | 0.2212 | 0.23 | -0.09 | 0.51 |
| Anxiety (STAI) | 63 | 35.78 | 10.44 | 63 | 32.11 | 7.11 | 1261.00 | 3.14 | 0.002 | 0.0047 | 0.47 | 0.22 | 0.67 |
| Arousal | 63 | 3.00 | 2.29 | 63 | 3.06 | 2.12 | 694.00 | 0.05 | 0.967 | 0.9670 | 0.01 | -0.30 | 0.31 |
| Valence | 63 | 7.52 | 1.61 | 63 | 7.48 | 2.34 | 574.00 | -0.38 | 0.703 | 0.8202 | -0.06 | -0.37 | 0.25 |
| POMS | 63 | 12.97 | 30.06 | 63 | 4.02 | 23.33 | 1326.50 | 3.33 | < .001 | 0.0004 | 0.50 | 0.25 | 0.69 |

**Table 1F.** Nonparametric t-test (Wilcoxon Signed-Rank) of timepoint (pre vs post) differences for the VR group. 'DV' refers to dependent variable. 'n' refers to group sample size. 'M' and 'SD' refer to mean and standard deviation. 'p.raw' and 'p.fdr' refer to the uncorrected p value and FDR-corrected p value. 'RBC' refers to rank biserial correlation as a measure of effect size (ranging from -1 to 1) and reflects the relative proportions of positive and negative ranks. 'RBC (low)' and 'RBC (high)' refer to the lower and upper bounds of the 95% confidence interval of RBC.

#### Table 1G (Breathwork)

| Group (Breathwork) | | | | | | | | | | | | | |
|---|---|---|---|---|---|---|---|---|---|---|---|---|---|
| | Pre | | | Post | | | Wilcoxon Signed-Rank test | | | | | | |
| **DV** | **n** | **M** | **SD** | **n** | **M** | **SD** | **W** | **Z** | **p.raw** | **p.fdr** | **RBC** | **RBC (low)** | **RBC (high)** |
| TMT (RTACC) | 63 | 167.01 | 152.12 | 61 | 145.42 | 125.07 | 1315.00 | 2.65 | 0.008 | 0.0140 | 0.39 | 0.12 | 0.61 |
| Architex (Total Speed) | 57 | 219.05 | 215.90 | 57 | 137.35 | 51.17 | 1480.00 | 5.19 | < .001 | 0.0002 | 0.79 | 0.65 | 0.88 |
| Architex (Total Score) | 57 | 38.70 | 3.25 | 57 | 38.04 | 3.46 | 870.00 | 1.94 | 0.052 | 0.0728 | 0.31 | 0.01 | 0.56 |
| Anxiety (STAI) | 64 | 36.06 | 9.87 | 64 | 31.34 | 8.05 | 1435.00 | 4.15 | < .001 | 0.0002 | 0.62 | 0.41 | 0.77 |
| Arousal | 64 | 2.58 | 1.67 | 64 | 2.84 | 1.85 | 519.00 | -0.71 | 0.478 | 0.4780 | -0.12 | -0.42 | 0.20 |
| Valence | 64 | 7.09 | 1.67 | 64 | 7.53 | 2.45 | 436.00 | -1.56 | 0.118 | 0.1377 | -0.26 | -0.53 | 0.06 |
| POMS | 64 | 12.63 | 26.81 | 64 | 2.38 | 23.00 | 1448.00 | 4.25 | < .001 | 0.0002 | 0.64 | 0.43 | 0.78 |

**Table 1G.** Nonparametric t-test (Wilcoxon Signed-Rank) of timepoint (pre vs post) differences for the Breathwork group. 'DV' refers to dependent variable. 'n' refers to group sample size. 'M' and 'SD' refer to mean and standard deviation. 'p.raw' and 'p.fdr' refer to the uncorrected p value and FDR-corrected p value. 'RBC' refers to rank biserial correlation as a measure of effect size (ranging from -1 to 1) and reflects the relative proportions of positive and negative ranks. 'RBC (low)' and 'RBC (high)' refer to the lower and upper bounds of the 95% confidence interval of RBC.

#### Table 1H (Rain)

| Group (Rain) | | | | | | | | | | | | | |
|---|---|---|---|---|---|---|---|---|---|---|---|---|---|
| | Pre | | | Post | | | Wilcoxon Signed-Rank test | | | | | | |
| **DV** | **n** | **M** | **SD** | **n** | **M** | **SD** | **W** | **Z** | **p.raw** | **p.fdr** | **RBC** | **RBC (low)** | **RBC (high)** |
| TMT (RTACC) | 61 | 165.74 | 162.97 | 61 | 126.65 | 73.95 | 1429.00 | 3.47 | < .001 | 0.0002 | 0.51 | 0.27 | 0.69 |

| | | | | | | | | | | | | | |
|---|---|---|---|---|---|---|---|---|---|---|---|---|---|
| Architex (Total Speed) | 58 | 203.36 | 84.57 | 58 | 132.23 | 57.66 | 1567.00 | 5.88 | < .001 | 0.0002 | 0.90 | 0.82 | 0.94 |
| Architex (Total Score) | 58 | 37.85 | 5.68 | 58 | 37.72 | 6.15 | 586.50 | -0.26 | 0.799 | 0.7990 | -0.04 | -0.35 | 0.27 |
| Anxiety (STAI) | 62 | 36.26 | 9.46 | 62 | 33.45 | 6.37 | 1156.50 | 2.62 | 0.009 | 0.0158 | 0.40 | 0.12 | 0.62 |
| Arousal | 62 | 3.27 | 2.47 | 62 | 2.77 | 1.95 | 686.50 | 1.30 | 0.192 | 0.2240 | 0.22 | -0.11 | 0.50 |
| Valence | 62 | 7.52 | 1.66 | 62 | 7.74 | 1.83 | 395.50 | -1.38 | 0.162 | 0.2240 | -0.24 | -0.52 | 0.10 |
| POMS | 62 | 14.36 | 26.73 | 62 | 5.47 | 19.87 | 1403.50 | 3.91 | < .001 | 0.0002 | 0.59 | 0.36 | 0.75 |

**Table 1H.** Nonparametric t-test (Wilcoxon Signed-Rank) of timepoint (pre vs post) differences for the Rain group. 'DV' refers to dependent variable. 'n' refers to group sample size. 'M' and 'SD' refer to mean and standard deviation. 'p.raw' and 'p.fdr' refer to the uncorrected p value and FDR-corrected p value. 'RBC' refers to rank biserial correlation as a measure of effect size (ranging from -1 to 1) and reflects the relative proportions of positive and negative ranks. 'RBC (low)' and 'RBC (high)' refer to the lower and upper bounds of the 95% confidence interval of RBC.

#### Table 1I (All)

| **Group (All)** | | | | | | | | | | | | | |
|---|---|---|---|---|---|---|---|---|---|---|---|---|---|
| | **Pre** | | | **Post** | | | **Wilcoxon Signed-Rank test** | | | | | | |
| **DV** | **n** | **M** | **SD** | **n** | **M** | **SD** | **W** | **Z** | **p.raw** | **p.fdr** | **RBC** | **RBC (low)** | **RBC (high)** |
| TMT (RTACC) | 124 | 166.38 | 156.91 | 122 | 136.03 | 102.75 | 5443.00 | 4.32 | < .001 | 0.0002 | 0.45 | 0.27 | 0.60 |
| Architex (Total Speed) | 115 | 211.14 | 162.89 | 115 | 134.76 | 54.36 | 6044.00 | 7.82 | < .001 | 0.0002 | 0.84 | 0.77 | 0.90 |
| Architex (Total Score) | 115 | 38.27 | 4.64 | 115 | 37.88 | 4.98 | 2886.00 | 1.24 | 0.214 | 0.2497 | 0.14 | -0.08 | 0.35 |
| Anxiety (STAI) | 126 | 36.16 | 9.63 | 126 | 32.38 | 7.32 | 5156.50 | 4.86 | < .001 | 0.0002 | 0.52 | 0.35 | 0.66 |
| Arousal | 126 | 2.92 | 2.12 | 126 | 2.81 | 1.89 | 2418.00 | 0.51 | 0.606 | 0.6060 | 0.06 | -0.17 | 0.28 |
| Valence | 126 | 7.30 | 1.67 | 126 | 7.64 | 2.16 | 1626.00 | -2.14 | 0.030 | 0.0420 | -0.26 | -0.46 | -0.03 |
| POMS | 126 | 13.48 | 26.68 | 126 | 3.90 | 21.49 | 5659.00 | 5.77 | < .001 | 0.0002 | 0.61 | 0.47 | 0.73 |

**Table 1I.** Nonparametric t-test (Wilcoxon Signed-Rank) of timepoint (pre vs post) differences for all groups combined. 'DV' refers to dependent variable. 'n' refers to group sample size. 'M' and 'SD' refer to mean and standard deviation. 'p.raw' and 'p.fdr' refer to the uncorrected p value and FDR-corrected p value. 'RBC' refers to rank biserial correlation as a measure of effect size (ranging from -1 to 1) and reflects the relative proportions of positive and negative ranks. 'RBC (low)' and 'RBC (high)' refer to the lower and upper bounds of the 95% confidence interval of RBC.
