## Supplementary material for "Contrasting cognitive, behavioral, and physiological responses to breathwork vs. naturalistic stimuli in reflective chamber and VR headset environments": S2 Appendix.pdf

### S2 Appendix: Transcription of 4-8 Breathwork Audio

The following is a transcription of the audio guidance provided during the "4-8 Breathwork" condition of the study. This audio was accompanied by dynamic lighting patterns synchronized with the breathing instructions.

[Speaker: Dr. Janelle MacAulay]

Welcome back to the Lumina Infinity Cube. I'm Dr. Janelle MacAuley and thank you for joining me for a guided mindfulness breathing exercise. As we've discussed, mindfulness training is an effective way to build mental strength, increase self-awareness, and train our minds to live more in the present moment.

What I mean by that is instead of spending much of our day distracted and mind wandering about the future or even the past, we can train our minds to focus in the moment to live our lives more on the play button instead of rewind or fast forward.

Maybe you found yourself thinking a lot about work while you're at home with your family. Or maybe you found yourself thinking a lot about home while you're at work. Have you ever read a page in a book, got to the bottom and then thought to yourself, "I do not remember what I just read"? Or maybe you've driven your car somewhere and couldn't remember what roads you took to your destination.

Well, our minds are fantastic at mental time travel, therefore we found ourselves unintentionally thinking about other things when we're trying to stay on task with what's going on right in front of us. Mindfulness training helps us live where our feet are planted, to be focused in the moment whether that's at work or at home or reading a book, it's an extremely difficult thing to do, which is why we must train our minds.

To begin today's session, we will do a mindful minute to cultivate a sense of presence and awareness in this space. Focus on your breathing and the light patterns around you as you strengthen your attention system to stay in the present moment. Now remember your mental push-ups if you find yourself distracted, it's okay - you're not doing it wrong. Simply acknowledge that thought that distracted you and bring your attention back to the present moment.

Ready to begin ? Good. Now take one big inhale, and exhale. And find a comfortable position with your shoulders back and your head lifted.

[Music background]

And begin…

[Light and music starts]

3:14

Welcome back. Now that we've built a sense of moment awareness, let's begin our breathwork training.

One of the most powerful things we can do for ourselves, is prepare to perform at our best inside moments we may encounter adversity and stress, is to trigger our relaxation response. Our mind and body are so connected, we can that through a breathing technique where we elongate our exhale. How does that mechanism work exactly? Well, when our ancestors encountered say… a saber-toothed tiger. It would trigger their sympathetic nervous system to go into fight-or-flight mode. In most circumstances, they choose flight. And once they found themselves out of danger, they would take a long nice exhale that signified, "Phew..!! I am safe. I can find calm."

Well, that same ability exists inside you and me. Now we have the same ancient brain. It's just living in a modern world, so we might trigger the same stress response of our sympathetic nervous system while driving in traffic or dealing with an unpleasant situation at work. Instead of running away from a saber-toothed tiger, we can also use a deep and long exhale to energize our parasympathetic system signifying safety and calm in those situations

[Music]

4:37

Today's exercise will train your body and mind that when we take a deep breath accompanied by an elongated exhale, we are signaling our brain that it is okay to relax. Each time you exhale during this exercise, I want you to focus on relaxing your shoulders and elongating your neck as you cultivate a sense of calm and relaxation.

[Music]

5:00

Ready to begin? As we start our practice today, I want you to find a slow and smooth cadence of your breathing. Inhale and exhale through your nose.

[Music]

For this practice, I'm going to ask you to inhale for a count of four, and hold your breath at the top for one count, and exhale for a count of eight. Now let’s take a deep inhale for a count of four.

One… two... three... four...

One… two... three... four...  Hold your breath at the top.

Now exhale two… three… four... five... six… seven... eight ... Again, let’s take a deep inhale.

Two... three... four... Hold your breath at the top. Now enjoy your exhale.

Two... three... four… five.. six... seven... eight... Again, let's take a deep breath and inhale for a count of four.

One… two... three... four...

Hold your breath at the top. Enjoy your exhale.

Two... three... four… five.. six... seven... eight...

One… two... three... four... Hold.

One… Two... three... four… five.. six... seven... eight...

One… two... three... four... Hold.

One… Two... three... four… five.. six... seven... eight...

One… two... three... four... Hold.

One… Two... three... four… five.. six... seven... eight...

Continue to focus on your breath

7:11

Focus on the count. Let go of distractions. Enjoy the sensations of your exhale. Embrace the feelings of calm and the sense of relaxation. Remember your mental push-ups. Remember the thought. Come back to your breath and the count.

One… two... three... four... Hold.

One… Two... three... four… five.. six... seven... eight...

One… two... three... four... Hold.

One… Two... three... four… five.. six... seven... eight...

One… two... three... four... Hold.

One… Two... three... four… five.. six... seven... eight...

One… two... three... four... Hold.

One… Two... three... four… five.. six... seven... eight...

As you take your final deep inhalation and deep exhalation. Begin to find the normal cadence of your breathing. Inhale. Exhale. Inhale. Exhale. Focus on your chosen sensation. Inhale. Exhale.

Notice how you feel right now. Maybe more focused, steady calm. Emphasizing the exhale is meant to stimulate our parasympathetic nervous system which is the calming counterpart to our stress-induced sympathetic nervous system. The next time you find yourself in a situation where your sympathetic nervous system kicks into overdrive and you feel almost out of control, be confident in the fact that you’ve trained your mind enough, so you can find calm and safety with an elongated exhale. The more you practice this skill set, the more you will set your mind and your body for success, and the more available it will be to you, and the moments you need it the most
