## Supplementary material for "Contrasting cognitive, behavioral, and physiological responses to breathwork vs. naturalistic stimuli in reflective chamber and VR headset environments": S2 Tables A-F.pdf

### S2: Group differences in timepoint effects

##### Table 2A (MindGym Breathwork vs MindGym Rain)

| | MindGym (Breathwork) | | | MindGym (Rain) | | | Mann-Whitney U test | | | | | |
|---|---|---|---|---|---|---|---|---|---|---|---|---|
| **DV** | **n** | **ΔM** | **SD** | **n** | **ΔM** | **SD** | **W** | **p.raw** | **p.fdr** | **RBC** | **RBC (low)** | **RBC (high)** |
| TMT (RTACC) | 30 | 2.68 | 157.82 | 31 | -42.64 | 95.63 | 514 | 0.487 | 0.713 | 0.11 | -0.18 | 0.38 |
| Architex (Total Speed) | 30 | -51.77 | 67.90 | 30 | -72.88 | 61.68 | 512 | 0.366 | 0.713 | 0.14 | -0.16 | 0.41 |
| Architex (Total Score) | 30 | -0.93 | 2.97 | 30 | 0.65 | 2.50 | 298.5 | 0.025 | 0.175 | -0.34 | -0.57 | -0.06 |
| Anxiety (STAI) | 31 | -3.68 | 9.35 | 32 | -4.09 | 8.65 | 475.5 | 0.783 | 0.783 | -0.04 | -0.32 | 0.24 |
| Arousal | 31 | -0.19 | 1.83 | 32 | -0.38 | 2.30 | 471.5 | 0.736 | 0.783 | -0.05 | -0.33 | 0.23 |
| Valence | 31 | 1.03 | 2.76 | 32 | 0.41 | 2.03 | 546 | 0.486 | 0.713 | 0.10 | -0.18 | 0.37 |
| POMS | 31 | -9.90 | 17.48 | 32 | -10.50 | 13.34 | 544.5 | 0.509 | 0.713 | 0.10 | -0.19 | 0.37 |

**Table 2A.** Nonparametric t-tests (Mann-Whitney U) between the MindGym (Breathwork) and MindGym (Rain) groups. 'DV' refers to dependent variable. 'n' refers to group sample size. 'ΔM' and 'SD' refer to mean and standard deviation of the difference score (post – pre). 'p.raw' and 'p.fdr' refer to the uncorrected p value and FDR-corrected p value. 'RBC' refers to rank biserial correlation as a measure of effect size (ranging from -1 to 1) and reflects the relative proportions of positive and negative ranks. 'RBC (low)' and 'RBC (high)' refer to the lower and upper bounds of the 95% confidence interval of RBC.

##### Table 2B (MindGym Breathwork vs VR Breathwork)

| | MindGym (Breathwork) | | | VR (Breathwork) | | | Mann-Whitney U test | | | | | |
|---|---|---|---|---|---|---|---|---|---|---|---|---|
| **DV** | **n** | **ΔM** | **SD** | **n** | **ΔM** | **SD** | **W** | **p.raw** | **p.fdr** | **RBC** | **RBC (low)** | **RBC (high)** |
| TMT (RTACC) | 30 | 2.68 | 157.82 | 31 | -35.06 | 176.44 | 416.00 | 0.487 | 0.852 | -0.11 | -0.38 | 0.18 |
| Architex (Total Speed) | 30 | -51.77 | 67.90 | 27 | -114.97 | 301.49 | 433.00 | 0.663 | 0.856 | 0.07 | -0.23 | 0.36 |
| Architex (Total Score) | 30 | -0.93 | 2.97 | 27 | -0.37 | 2.55 | 357.00 | 0.446 | 0.852 | -0.12 | -0.40 | 0.18 |
| Anxiety (STAI) | 31 | -3.68 | 9.35 | 33 | -5.70 | 8.41 | 534.00 | 0.767 | 0.856 | 0.04 | -0.24 | 0.32 |
| Arousal | 31 | -0.19 | 1.83 | 33 | 0.70 | 2.76 | 417.50 | 0.203 | 0.852 | -0.18 | -0.44 | 0.10 |
| Valence | 31 | 1.03 | 2.76 | 33 | -0.12 | 3.23 | 582.00 | 0.341 | 0.852 | 0.14 | -0.15 | 0.40 |
| POMS | 31 | -9.90 | 17.48 | 33 | -10.58 | 20.44 | 525.50 | 0.856 | 0.856 | 0.03 | -0.25 | 0.30 |

**Table 2B.** Nonparametric t-tests (Mann-Whitney U) between the MindGym (Breathwork) and VR (Breathwork) groups. 'DV' refers to dependent variable. 'n' refers to group sample size. 'ΔM' and 'SD' refer to mean and standard deviation of the difference score (post – pre). 'p.raw' and 'p.fdr' refer to the uncorrected p value and FDR-corrected p value. 'RBC' refers to rank biserial correlation as a measure of effect size (ranging from -1 to 1) and reflects the relative proportions of positive and negative ranks. 'RBC (low)' and 'RBC (high)' refer to the lower and upper bounds of the 95% confidence interval of RBC.

##### Table 2C (MindGym Rain vs VR Rain)

| | MindGym (Rain) | | | VR (Rain) | | | Mann-Whitney U test | | | | | |
|---|---|---|---|---|---|---|---|---|---|---|---|---|
| **DV** | **n** | **ΔM** | **SD** | **n** | **ΔM** | **SD** | **W** | **p.raw** | **p.fdr** | **RBC** | **RBC (low)** | **RBC (high)** |
| TMT (RTACC) | 31 | -42.64 | 95.63 | 30 | -35.42 | 206.73 | 347.00 | 0.090 | 0.214 | -0.25 | -0.50 | 0.03 |
| Architex (Total Speed) | 30 | -72.88 | 61.68 | 28 | -69.26 | 73.95 | 409.00 | 0.871 | 0.871 | -0.03 | -0.32 | 0.27 |
| Architex (Total Score) | 30 | 0.65 | 2.50 | 28 | -0.96 | 3.74 | 519.50 | 0.122 | 0.214 | 0.24 | -0.06 | 0.49 |
| Anxiety (STAI) | 32 | -4.09 | 8.65 | 30 | -1.43 | 7.17 | 367.50 | 0.114 | 0.214 | -0.23 | -0.48 | 0.05 |

| | | | | | | | | | | | |
|---|---|---|---|---|---|---|---|---|---|---|---|
| Arousal | 32 | -0.38 | 2.30 | 30 | -0.63 | 2.87 | 521.50 | 0.557 | 0.650 | 0.09 | -0.20 | 0.36 |
| Valence | 32 | 0.41 | 2.03 | 30 | 0.03 | 1.81 | 559.50 | 0.256 | 0.358 | 0.17 | -0.12 | 0.43 |
| POMS | 32 | -10.50 | 13.34 | 30 | -7.17 | 19.80 | 350.50 | 0.069 | 0.214 | -0.27 | -0.51 | 0.01 |

**Table 2C.** Nonparametric t-tests (Mann-Whitney U) between the MindGym (Rain) and VR (Rain) groups. 'DV' refers to dependent variable. 'n' refers to group sample size. 'ΔM' and 'SD' refer to mean and standard deviation of the difference score (post – pre). 'p.raw' and 'p.fdr' refer to the uncorrected p value and FDR-corrected p value. 'RBC' refers to rank biserial correlation as a measure of effect size (ranging from -1 to 1) and reflects the relative proportions of positive and negative ranks. 'RBC (low)' and 'RBC (high)' refer to the lower and upper bounds of the 95% confidence interval of RBC.

#### Table 2D (VR Breathwork vs VR Rain)

| | VR (Breathwork) | | | VR (Rain) | | | Mann-Whitney U test | | | | | |
|---|---|---|---|---|---|---|---|---|---|---|---|---|
| **DV** | **n** | **ΔM** | **SD** | **n** | **ΔM** | **SD** | **W** | **p.raw** | **p.fdr** | **RBC** | **RBC (low)** | **RBC (high)** |
| TMT (RTACC) | 31 | -35.06 | 176.44 | 30 | -35.42 | 206.73 | 456.00 | 0.903 | 0.939 | -0.02 | -0.30 | 0.27 |
| Architex (Total Speed) | 27 | -114.97 | 301.49 | 28 | -69.26 | 73.95 | 395.00 | 0.783 | 0.939 | 0.05 | -0.26 | 0.34 |
| Architex (Total Score) | 27 | -0.37 | 2.55 | 28 | -0.96 | 3.74 | 373.00 | 0.939 | 0.939 | -0.01 | -0.31 | 0.29 |
| Anxiety (STAI) | 33 | -5.70 | 8.41 | 30 | -1.43 | 7.17 | 347.00 | 0.042 | 0.294 | -0.30 | -0.53 | -0.02 |
| Arousal | 33 | 0.70 | 2.76 | 30 | -0.63 | 2.87 | 610.50 | 0.110 | 0.385 | 0.23 | -0.05 | 0.48 |
| Valence | 33 | -0.12 | 3.23 | 30 | 0.03 | 1.81 | 533.00 | 0.601 | 0.939 | 0.08 | -0.21 | 0.35 |
| POMS | 33 | -10.58 | 20.44 | 30 | -7.17 | 19.80 | 408.50 | 0.236 | 0.551 | -0.18 | -0.43 | 0.11 |

**Table 2D.** Nonparametric t-tests (Mann-Whitney U) between the VR (Breathwork) and VR (Rain) groups. 'DV' refers to dependent variable. 'n' refers to group sample size. 'ΔM' and 'SD' refer to mean and standard deviation of the difference score (post – pre). 'p.raw' and 'p.fdr' refer to the uncorrected p value and FDR-corrected p value. 'RBC' refers to rank biserial correlation as a measure of effect size (ranging from -1 to 1) and reflects the relative proportions of positive and negative ranks. 'RBC (low)' and 'RBC (high)' refer to the lower and upper bounds of the 95% confidence interval of RBC.

#### Table 2E (Breathwork vs Rain)

| | Breathwork | | | Rain | | | Mann-Whitney U test | | | | | |
|---|---|---|---|---|---|---|---|---|---|---|---|---|
| **DV** | **n** | **ΔM** | **SD** | **n** | **ΔM** | **SD** | **W** | **p.raw** | **p.fdr** | **RBC** | **RBC (low)** | **RBC (high)** |
| TMT (RTACC) | 61 | -16.50 | 167.23 | 61 | -39.09 | 158.87 | 1936.00 | 0.701 | 0.705 | 0.04 | -0.16 | 0.24 |
| Architex (Total Speed) | 57 | -81.71 | 213.55 | 58 | -71.13 | 67.30 | 1804.00 | 0.400 | 0.560 | 0.09 | -0.12 | 0.30 |
| Architex (Total Score) | 57 | -0.67 | 2.77 | 58 | -0.13 | 3.24 | 1353.50 | 0.094 | 0.480 | -0.18 | -0.38 | 0.03 |
| Anxiety (STAI) | 64 | -4.72 | 8.86 | 62 | -2.81 | 8.02 | 1679.00 | 0.137 | 0.480 | -0.15 | -0.34 | 0.05 |
| Arousal | 64 | 0.27 | 2.38 | 62 | -0.50 | 2.57 | 2192.50 | 0.304 | 0.560 | 0.11 | -0.10 | 0.30 |
| Valence | 64 | 0.44 | 3.04 | 62 | 0.23 | 1.92 | 2162.00 | 0.379 | 0.560 | 0.09 | -0.11 | 0.28 |
| POMS | 64 | -10.25 | 18.92 | 62 | -8.89 | 16.73 | 1906.00 | 0.705 | 0.705 | -0.04 | -0.24 | 0.16 |

**Table 2E.** Nonparametric t-tests (Mann-Whitney U) between the Breathwork and Rain groups. 'DV' refers to dependent variable. 'n' refers to group sample size. 'ΔM' and 'SD' refer to mean and standard deviation of the difference score (post – pre). 'p.raw' and 'p.fdr' refer to the uncorrected p value and FDR-corrected p value. 'RBC' refers to rank biserial correlation as a measure of effect size (ranging from -1 to 1) and reflects the relative proportions of positive and negative ranks. 'RBC (low)' and 'RBC (high)' refer to the lower and upper bounds of the 95% confidence interval of RBC.

#### Table 2F (MindGym vs VR)

| | MindGym | | | VR | | | Mann-Whitney U test | | | | | |
|---|---|---|---|---|---|---|---|---|---|---|---|---|
| **DV** | **n** | **ΔM** | **SD** | **n** | **ΔM** | **SD** | **W** | **p.raw** | **p.fdr** | **RBC** | **RBC (low)** | **RBC (high)** |
| TMT (RTACC) | 61 | -20.35 | 130.89 | 61 | -35.24 | 190.32 | 1534.00 | 0.095 | 0.585 | -0.18 | -0.37 | 0.03 |
| Architex (Total Speed) | 60 | -62.33 | 65.19 | 55 | -91.70 | 216.87 | 1683.00 | 0.856 | 0.856 | 0.02 | -0.19 | 0.23 |
| Architex (Total Score) | 60 | -0.14 | 2.84 | 55 | -0.67 | 3.20 | 1778.50 | 0.472 | 0.661 | 0.08 | -0.13 | 0.28 |
| Anxiety (STAI) | 63 | -3.89 | 8.93 | 63 | -3.67 | 8.07 | 1829.00 | 0.449 | 0.661 | -0.08 | -0.27 | 0.12 |
| Arousal | 63 | -0.29 | 2.07 | 63 | 0.06 | 2.87 | 1875.50 | 0.592 | 0.691 | -0.06 | -0.25 | 0.15 |
| Valence | 63 | 0.71 | 2.42 | 63 | -0.05 | 2.63 | 2263.50 | 0.167 | 0.585 | 0.14 | -0.06 | 0.33 |
| POMS | 63 | -10.21 | 15.40 | 63 | -8.95 | 20.05 | 1772.00 | 0.301 | 0.661 | -0.11 | -0.30 | 0.10 |

**Table 2F.** Nonparametric t-tests (Mann-Whitney U) between the MindGym and VR groups. 'DV' refers to dependent variable. 'n' refers to group sample size. 'ΔM' and 'SD' refer to mean and standard deviation of the difference score (post – pre). 'p.raw' and 'p.fdr' refer to the uncorrected p value and FDR-corrected p value. 'RBC' refers to rank biserial correlation as a measure of effect size (ranging from -1 to 1) and reflects the relative proportions of positive and negative ranks. 'RBC (low)' and 'RBC (high)' refer to the lower and upper bounds of the 95% confidence interval of RBC.
