## Supplementary material for "Contrasting cognitive, behavioral, and physiological responses to breathwork vs. naturalistic stimuli in reflective chamber and VR headset environments": S3 Tables A-F.pdf

### S3: Group differences in Post variables

##### Table 3A (MindGym Breathwork vs MindGym Rain)

| | MindGym (Breathwork) | | | MindGym (Rain) | | | Mann-Whitney U test | | | | | |
|---|---|---|---|---|---|---|---|---|---|---|---|---|
| **DV** | **n** | **M** | **SD** | **n** | **M** | **SD** | **W** | **p.raw** | **p.fdr** | **RBC** | **RBC (low)** | **RBC (high)** |
| Awe | 31 | 118.48 | 34.24 | 32 | 113.94 | 35.23 | 543.50 | 0.518 | 0.637 | 0.10 | -0.27 | 0.44 |
| BSE | 31 | 4.16 | 1.64 | 32 | 4.28 | 1.73 | 477.00 | 0.795 | 0.795 | -0.04 | -0.39 | 0.33 |
| EDI | 28 | 4.21 | 2.11 | 30 | 3.54 | 2.40 | 505.00 | 0.189 | 0.548 | 0.20 | -0.19 | 0.54 |
| Immersion | 31 | 67.23 | 11.95 | 32 | 62.78 | 15.21 | 572.00 | 0.299 | 0.548 | 0.15 | -0.22 | 0.49 |
| IPQ (General) | 31 | 0.68 | 1.72 | 32 | 0.91 | 1.73 | 450.00 | 0.521 | 0.637 | -0.09 | -0.44 | 0.28 |
| IPQ (Spatial Presence) | 31 | 3.03 | 3.64 | 32 | 2.06 | 3.56 | 582.50 | 0.234 | 0.548 | 0.17 | -0.20 | 0.50 |
| IPQ (Involvement) | 31 | 4.16 | 3.45 | 32 | 1.97 | 4.13 | 633.00 | 0.059 | 0.336 | 0.28 | -0.09 | 0.58 |
| IPQ (Experience Realism) | 31 | -2.68 | 3.26 | 32 | -1.91 | 3.75 | 417.50 | 0.280 | 0.548 | -0.16 | -0.49 | 0.22 |
| Motion (Overall) | 31 | 19.00 | 10.18 | 32 | 17.80 | 10.69 | 527.00 | 0.674 | 0.741 | 0.06 | -0.31 | 0.41 |
| Toronto (Curiosity) | 31 | 13.84 | 5.79 | 32 | 16.59 | 4.26 | 359.50 | 0.061 | 0.336 | -0.28 | -0.58 | 0.10 |
| Toronto (Decentering) | 31 | 16.65 | 5.96 | 32 | 16.09 | 4.04 | 549.00 | 0.469 | 0.637 | 0.11 | -0.26 | 0.45 |

**Table 3A.** Nonparametric t-tests (Mann-Whitney U) between the MindGym (Breathwork) and MindGym (Rain) groups. 'DV' refers to dependent variable. 'n' refers to group sample size. 'M' and 'SD' refer to mean and standard deviation. 'p.raw' and 'p.fdr' refer to the uncorrected p value and FDR-corrected p value. 'RBC' refers to rank biserial correlation as a measure of effect size (ranging from -1 to 1) and reflects the relative proportions of positive and negative ranks. 'RBC (low)' and 'RBC (high)' refer to the lower and upper bounds of the 95% confidence interval of RBC.

##### Table 3B (MindGym Breathwork vs VR Breathwork)

| | MindGym (Breathwork) | | | VR (Breathwork) | | | Mann-Whitney U test | | | | | |
|---|---|---|---|---|---|---|---|---|---|---|---|---|
| **DV** | **n** | **M** | **SD** | **n** | **M** | **SD** | **W** | **p.raw** | **p.fdr** | **RBC** | **RBC (low)** | **RBC (high)** |
| Awe | 31 | 118.48 | 34.24 | 33 | 105.76 | 34.46 | 632.00 | 0.107 | 0.782 | 0.24 | -0.13 | 0.55 |
| BSE | 31 | 4.16 | 1.64 | 33 | 3.91 | 1.51 | 553.50 | 0.569 | 0.782 | 0.08 | -0.28 | 0.43 |
| EDI | 28 | 4.21 | 2.11 | 27 | 3.89 | 2.24 | 414.50 | 0.544 | 0.782 | 0.10 | -0.30 | 0.46 |
| Immersion | 31 | 67.23 | 11.95 | 33 | 64.21 | 12.61 | 574.00 | 0.405 | 0.782 | 0.12 | -0.25 | 0.46 |
| IPQ (General) | 31 | 0.68 | 1.72 | 33 | 0.97 | 1.81 | 463.00 | 0.510 | 0.782 | -0.10 | -0.44 | 0.27 |
| IPQ (Spatial Presence) | 31 | 3.03 | 3.64 | 33 | 3.06 | 3.17 | 517.00 | 0.946 | 0.946 | 0.00 | -0.36 | 0.36 |
| IPQ (Involvement) | 31 | 4.16 | 3.45 | 33 | 3.15 | 4.19 | 582.50 | 0.341 | 0.782 | 0.14 | -0.23 | 0.47 |
| IPQ (Experience Realism) | 31 | -2.68 | 3.26 | 33 | -2.61 | 3.31 | 544.50 | 0.660 | 0.807 | 0.07 | -0.30 | 0.41 |
| Motion (Overall) | 31 | 19.00 | 10.18 | 33 | 16.81 | 11.08 | 600.50 | 0.232 | 0.782 | 0.17 | -0.20 | 0.50 |
| Toronto (Curiosity) | 31 | 13.84 | 5.79 | 33 | 14.88 | 6.31 | 464.50 | 0.532 | 0.782 | -0.09 | -0.44 | 0.28 |
| Toronto (Decentering) | 31 | 16.65 | 5.96 | 33 | 17.18 | 4.48 | 506.00 | 0.946 | 0.946 | -0.01 | -0.37 | 0.35 |

**Table 3B.** Nonparametric t-tests (Mann-Whitney U) between the MindGym (Breathwork) and VR (Breathwork) groups. 'DV' refers to dependent variable. 'n' refers to group sample size. 'M' and 'SD' refer to mean and standard deviation. 'p.raw' and 'p.fdr' refer to the uncorrected p value and FDR-corrected p value. 'RBC' refers to rank biserial correlation as a measure of effect size (ranging from -1 to 1) and reflects the relative proportions of positive and negative ranks. 'RBC (low)' and 'RBC (high)' refer to the lower and upper bounds of the 95% confidence interval of RBC.

#### Table 3C (MindGym Rain vs VR Rain)

| | MindGym (Rain) | | | VR (Rain) | | | Mann-Whitney U test | | | | | |
|---|---|---|---|---|---|---|---|---|---|---|---|---|
| **DV** | **n** | **M** | **SD** | **n** | **M** | **SD** | **W** | **p.raw** | **p.fdr** | **RBC** | **RBC (low)** | **RBC (high)** |
| Awe | 32 | 113.94 | 35.23 | 30 | 114.83 | 29.69 | 478.50 | 0.989 | 0.989 | 0.00 | -0.37 | 0.36 |
| BSE | 32 | 4.28 | 1.73 | 30 | 4.10 | 1.67 | 515.00 | 0.620 | 0.950 | 0.07 | -0.30 | 0.43 |
| EDI | 30 | 3.54 | 2.40 | 27 | 3.35 | 2.67 | 430.50 | 0.689 | 0.950 | 0.06 | -0.32 | 0.43 |
| Immersion | 32 | 62.78 | 15.21 | 30 | 61.97 | 10.71 | 525.00 | 0.530 | 0.950 | 0.09 | -0.28 | 0.44 |
| IPQ (General) | 32 | 0.91 | 1.73 | 30 | 1.17 | 1.34 | 453.50 | 0.706 | 0.950 | -0.06 | -0.41 | 0.32 |
| IPQ (Spatial Presence) | 32 | 2.06 | 3.56 | 30 | 3.23 | 3.77 | 380.50 | 0.161 | 0.886 | -0.21 | -0.53 | 0.17 |
| IPQ (Involvement) | 32 | 1.97 | 4.13 | 30 | 1.33 | 4.11 | 559.00 | 0.267 | 0.950 | 0.17 | -0.21 | 0.50 |
| IPQ (Experience Realism) | 32 | -1.91 | 3.75 | 30 | -3.13 | 2.97 | 629.00 | 0.035 | 0.385 | 0.31 | -0.06 | 0.61 |
| Motion (Overall) | 32 | 17.80 | 10.69 | 30 | 16.83 | 6.68 | 473.00 | 0.927 | 0.989 | -0.02 | -0.38 | 0.35 |
| Toronto (Curiosity) | 32 | 16.59 | 4.26 | 30 | 16.60 | 6.32 | 459.50 | 0.777 | 0.950 | -0.04 | -0.40 | 0.33 |
| Toronto (Decentering) | 32 | 16.09 | 4.04 | 30 | 16.87 | 5.79 | 423.00 | 0.424 | 0.950 | -0.12 | -0.46 | 0.26 |

**Table 3C.** Nonparametric t-tests (Mann-Whitney U) between the MindGym (Rain) and VR (Rain) groups. 'DV' refers to dependent variable. 'n' refers to group sample size. 'M' and 'SD' refer to mean and standard deviation. 'p.raw' and 'p.fdr' refer to the uncorrected p value and FDR-corrected p value. 'RBC' refers to rank biserial correlation as a measure of effect size (ranging from -1 to 1) and reflects the relative proportions of positive and negative ranks. 'RBC (low)' and 'RBC (high)' refer to the lower and upper bounds of the 95% confidence interval of RBC.

#### Table 3D (VR Breathwork vs VR Rain)

| | VR (Breathwork) | | | VR (Rain) | | | Mann-Whitney U test | | | | | |
|---|---|---|---|---|---|---|---|---|---|---|---|---|
| **DV** | **n** | **M** | **SD** | **n** | **M** | **SD** | **W** | **p.raw** | **p.fdr** | **RBC** | **RBC (low)** | **RBC (high)** |
| Awe | 33 | 105.76 | 34.46 | 30 | 114.83 | 29.69 | 408.00 | 0.234 | 0.673 | -0.18 | -0.51 | 0.20 |
| BSE | 33 | 3.91 | 1.51 | 30 | 4.10 | 1.67 | 467.00 | 0.699 | 0.879 | -0.06 | -0.41 | 0.31 |
| EDI | 27 | 3.89 | 2.24 | 27 | 3.35 | 2.67 | 406.00 | 0.478 | 0.751 | 0.11 | -0.29 | 0.48 |
| Immersion | 33 | 64.21 | 12.61 | 30 | 61.97 | 10.71 | 561.00 | 0.367 | 0.673 | 0.13 | -0.24 | 0.47 |
| IPQ (General) | 33 | 0.97 | 1.81 | 30 | 1.17 | 1.34 | 479.50 | 0.832 | 0.915 | -0.03 | -0.39 | 0.33 |
| IPQ (Spatial Presence) | 33 | 3.06 | 3.17 | 30 | 3.23 | 3.77 | 468.50 | 0.719 | 0.879 | -0.05 | -0.41 | 0.31 |
| IPQ (Involvement) | 33 | 3.15 | 4.19 | 30 | 1.33 | 4.11 | 620.00 | 0.086 | 0.673 | 0.25 | -0.12 | 0.56 |
| IPQ (Experience Realism) | 33 | -2.61 | 3.31 | 30 | -3.13 | 2.97 | 564.00 | 0.343 | 0.673 | 0.14 | -0.23 | 0.48 |
| Motion (Overall) | 33 | 16.81 | 11.08 | 30 | 16.83 | 6.68 | 421.50 | 0.313 | 0.673 | -0.15 | -0.48 | 0.23 |
| Toronto (Curiosity) | 33 | 14.88 | 6.31 | 30 | 16.60 | 6.32 | 414.00 | 0.267 | 0.673 | -0.16 | -0.50 | 0.21 |
| Toronto (Decentering) | 33 | 17.18 | 4.48 | 30 | 16.87 | 5.79 | 493.50 | 0.989 | 0.989 | 0.00 | -0.36 | 0.36 |

**Table 3D.** Nonparametric t-tests (Mann-Whitney U) between the VR (Breathwork) and VR (Rain) groups. 'DV' refers to dependent variable. 'n' refers to group sample size. 'M' and 'SD' refer to mean and standard deviation. 'p.raw' and 'p.fdr' refer to the uncorrected p value and FDR-corrected p value. 'RBC' refers to rank biserial correlation as a measure of effect size (ranging from -1 to 1) and reflects the relative proportions of positive and negative ranks. 'RBC (low)' and 'RBC (high)' refer to the lower and upper bounds of the 95% confidence interval of RBC.

#### Table 3E (Breathwork vs Rain)

| | Breathwork | | | Rain | | | Mann-Whitney U test | | | | | |
|---|---|---|---|---|---|---|---|---|---|---|---|---|
| **DV** | **n** | **M** | **SD** | **n** | **M** | **SD** | **W** | **p.raw** | **p.fdr** | **RBC** | **RBC (low)** | **RBC (high)** |
| Awe | 64 | 111.92 | 34.68 | 62 | 114.37 | 32.41 | 1928.50 | 0.788 | 0.854 | -0.03 | -0.29 | 0.23 |
| BSE | 64 | 4.03 | 1.56 | 62 | 4.19 | 1.69 | 1877.50 | 0.598 | 0.787 | -0.05 | -0.31 | 0.21 |
| EDI | 55 | 4.05 | 2.16 | 57 | 3.45 | 2.51 | 1818.00 | 0.146 | 0.501 | 0.16 | -0.12 | 0.42 |
| Immersion | 64 | 65.67 | 12.29 | 62 | 62.39 | 13.12 | 2258.00 | 0.182 | 0.501 | 0.14 | -0.13 | 0.38 |
| IPQ (General) | 64 | 0.83 | 1.76 | 62 | 1.03 | 1.55 | 1859.50 | 0.535 | 0.787 | -0.06 | -0.32 | 0.20 |
| IPQ (Spatial Presence) | 64 | 3.05 | 3.38 | 62 | 2.63 | 3.68 | 2115.00 | 0.522 | 0.787 | 0.07 | -0.20 | 0.32 |
| IPQ (Involvement) | 64 | 3.64 | 3.85 | 62 | 1.66 | 4.10 | 2501.50 | 0.011 | 0.121 | 0.26 | 0.00 | 0.49 |
| IPQ (Experience Realism) | 64 | -2.64 | 3.26 | 62 | -2.50 | 3.42 | 1946.00 | 0.854 | 0.854 | -0.02 | -0.28 | 0.24 |
| Motion (Overall) | 64 | 17.87 | 10.62 | 62 | 17.33 | 8.92 | 1889.00 | 0.644 | 0.787 | -0.05 | -0.30 | 0.22 |
| Toronto (Curiosity) | 64 | 14.38 | 6.04 | 62 | 16.60 | 5.31 | 1582.00 | 0.050 | 0.275 | -0.20 | -0.44 | 0.06 |
| Toronto (Decentering) | 64 | 16.92 | 5.22 | 62 | 16.47 | 4.94 | 2096.50 | 0.584 | 0.787 | 0.06 | -0.21 | 0.31 |

**Table 3E.** Nonparametric t-tests (Mann-Whitney U) between the Breathwork and Rain groups. ‘DV’ refers to dependent variable. ‘n’ refers to group sample size. ‘M’ and ‘SD’ refer to mean and standard deviation. ‘p.raw’ and ‘p.fdr’ refer to the uncorrected p value and FDR-corrected p value. ‘RBC’ refers to rank biserial correlation as a measure of effect size (ranging from -1 to 1) and reflects the relative proportions of positive and negative ranks. ‘RBC (low)’ and ‘RBC (high)’ refer to the lower and upper bounds of the 95% confidence interval of RBC.

#### Table 3F (MindGym vs VR)

| | MindGym | | | VR | | | Mann-Whitney U test | | | | | |
|---|---|---|---|---|---|---|---|---|---|---|---|---|
| **DV** | **n** | **M** | **SD** | **n** | **M** | **SD** | **W** | **p.raw** | **p.fdr** | **RBC** | **RBC (low)** | **RBC (high)** |
| Awe | 63 | 116.18 | 34.54 | 63 | 110.08 | 32.34 | 2236.50 | 0.220 | 0.586 | 0.13 | -0.14 | 0.38 |
| BSE | 63 | 4.22 | 1.67 | 63 | 4.00 | 1.58 | 2140.00 | 0.440 | 0.586 | 0.08 | -0.19 | 0.33 |
| EDI | 58 | 3.86 | 2.27 | 54 | 3.62 | 2.45 | 1664.00 | 0.570 | 0.586 | 0.06 | -0.22 | 0.33 |
| Immersion | 63 | 64.97 | 13.78 | 63 | 63.14 | 11.71 | 2210.00 | 0.272 | 0.586 | 0.11 | -0.15 | 0.36 |
| IPQ (General) | 63 | 0.79 | 1.72 | 63 | 1.06 | 1.60 | 1819.50 | 0.410 | 0.586 | -0.08 | -0.34 | 0.18 |
| IPQ (Spatial Presence) | 63 | 2.54 | 3.60 | 63 | 3.14 | 3.44 | 1783.50 | 0.326 | 0.586 | -0.10 | -0.35 | 0.16 |
| IPQ (Involvement) | 63 | 3.05 | 3.94 | 63 | 2.29 | 4.22 | 2250.00 | 0.194 | 0.586 | 0.13 | -0.13 | 0.38 |
| IPQ (Experience Realism) | 63 | -2.29 | 3.51 | 63 | -2.86 | 3.14 | 2359.00 | 0.067 | 0.586 | 0.19 | -0.08 | 0.43 |
| Motion (Overall) | 63 | 18.39 | 10.37 | 63 | 16.82 | 9.17 | 2162.00 | 0.386 | 0.586 | 0.09 | -0.18 | 0.34 |
| Toronto (Curiosity) | 63 | 15.24 | 5.21 | 63 | 15.70 | 6.33 | 1872.50 | 0.586 | 0.586 | -0.06 | -0.31 | 0.21 |
| Toronto (Decentering) | 63 | 16.37 | 5.05 | 63 | 17.03 | 5.11 | 1851.00 | 0.515 | 0.586 | -0.07 | -0.32 | 0.20 |

**Table 3F.** Nonparametric t-tests (Mann-Whitney U) between the MindGym and VR groups. ‘DV’ refers to dependent variable. ‘n’ refers to group sample size. ‘M’ and ‘SD’ refer to mean and standard deviation. ‘p.raw’ and ‘p.fdr’ refer to the uncorrected p value and FDR-corrected p value. ‘RBC’ refers to rank biserial correlation as a measure of effect size (ranging from -1 to 1) and reflects the relative proportions of positive and negative ranks. ‘RBC (low)’ and ‘RBC (high)’ refer to the lower and upper bounds of the 95% confidence interval of RBC.
