## Supplementary material for "Contrasting cognitive, behavioral, and physiological responses to breathwork vs. naturalistic stimuli in reflective chamber and VR headset environments": S4 Tables A-E.pdf

### S4: Group differences in Physio variables

#### Table 4A (MindGym Breathwork vs MindGym Rain)

| | MindGym (Breathwork) | | | MindGym (Rain) | | | Mann-Whitney U test | | | | | |
|---|---|---|---|---|---|---|---|---|---|---|---|---|
| DV | n | M | SD | n | M | SD | W | p.raw | p.fdr | RBC | RBC (low) | RBC (high) |
| Breath Rate (first 2 min) | 27 | 19.52 | 3.94 | 18 | 16.22 | 3.07 | 366.50 | 0.004 | 0.012 | 0.51 | 0.21 | 0.72 |
| Breath Rate (last 2 min) | 27 | 12.89 | 3.06 | 18 | 16.67 | 2.18 | 77.00 | < .001 | < .001 | -0.68 | -0.83 | -0.45 |
| Breath Rate change (last - first) | 27 | -6.63 | 5.59 | 18 | 0.44 | 3.11 | 68.50 | < .001 | < .001 | -0.72 | -0.85 | -0.51 |
| EDA (first 2 min) | 26 | 32.87 | 9.95 | 25 | 41.05 | 8.62 | 166.00 | 0.002 | 0.008 | -0.49 | -0.69 | -0.21 |
| EDA (last 2 min) | 15 | 38.22 | 11.86 | 12 | 46.43 | 8.04 | 56.00 | 0.103 | 0.247 | -0.38 | -0.69 | 0.05 |
| EDA change (last - first) | 15 | 4.91 | 8.82 | 12 | 6.82 | 5.78 | 75.00 | 0.486 | 0.667 | -0.17 | -0.55 | 0.27 |
| HR (first 1 min) | 20 | 67.78 | 10.44 | 14 | 70.17 | 12.95 | 124.00 | 0.592 | 0.710 | -0.11 | -0.47 | 0.28 |
| HR (last 1 min) | 20 | 70.91 | 9.29 | 14 | 72.03 | 12.80 | 134.00 | 0.849 | 0.926 | -0.04 | -0.42 | 0.34 |
| HR change (last - first) | 20 | 3.13 | 5.32 | 14 | 1.86 | 4.26 | 172.00 | 0.274 | 0.548 | 0.23 | -0.17 | 0.56 |
| HRV (first 5 min) | 20 | 0.21 | 0.04 | 14 | 0.20 | 0.04 | 165.00 | 0.396 | 0.667 | 0.18 | -0.22 | 0.52 |
| HRV (last 5 min) | 20 | 0.24 | 0.11 | 14 | 0.20 | 0.07 | 160.00 | 0.500 | 0.667 | 0.14 | -0.25 | 0.50 |
| HRV change (last - first) | 20 | 0.03 | 0.10 | 14 | 0.01 | 0.07 | 142.00 | 0.959 | 0.959 | 0.01 | -0.37 | 0.39 |

**Table 4A.** Nonparametric t-tests (Mann-Whitney U) between the MindGym (Breathwork) and MindGym (Rain) groups. 'DV' refers to dependent variable. 'n' refers to group sample size. 'M' and 'SD' refer to mean and standard deviation. 'p.raw' and 'p.fdr' refer to the uncorrected p value and FDR-corrected p value. 'RBC' refers to rank biserial correlation as a measure of effect size (ranging from -1 to 1) and reflects the relative proportions of positive and negative ranks. 'RBC (low)' and 'RBC (high)' refer to the lower and upper bounds of the 95% confidence interval of RBC.

#### Table 4B (MindGym Breathwork vs VR Breathwork)

| | MindGym (Breathwork) | | | VR (Breathwork) | | | Mann-Whitney U test | | | | | |
|---|---|---|---|---|---|---|---|---|---|---|---|---|
| DV | n | M | SD | n | M | SD | W | p.raw | p.fdr | RBC | RBC (low) | RBC (high) |
| Breath Rate (first 2 min) | 27 | 19.52 | 3.94 | 25 | 17.74 | 3.57 | 402.00 | 0.240 | 0.552 | 0.19 | -0.12 | 0.47 |
| Breath Rate (last 2 min) | 27 | 12.89 | 3.06 | 25 | 12.06 | 3.53 | 393.50 | 0.308 | 0.552 | 0.17 | -0.15 | 0.45 |
| Breath Rate change (last - first) | 27 | -6.63 | 5.59 | 25 | -5.68 | 4.52 | 307.00 | 0.582 | 0.657 | -0.09 | -0.39 | 0.22 |
| EDA (first 2 min) | 26 | 32.87 | 9.95 | 21 | 39.51 | 12.05 | 173.00 | 0.032 | 0.192 | -0.37 | -0.62 | -0.05 |
| EDA (last 2 min) | 15 | 38.22 | 11.86 | 21 | 41.75 | 11.90 | 130.00 | 0.391 | 0.552 | -0.18 | -0.51 | 0.21 |
| EDA change (last - first) | 15 | 4.91 | 8.82 | 21 | 2.24 | 5.55 | 187.00 | 0.357 | 0.552 | 0.19 | -0.20 | 0.52 |
| HR (first 1 min) | 20 | 67.78 | 10.44 | 20 | 71.79 | 11.24 | 166.00 | 0.369 | 0.552 | -0.17 | -0.49 | 0.19 |

| | | | | | | | | | | | | |
|---|---|---|---|---|---|---|---|---|---|---|---|---|
| HR (last 1 min) | 20 | 70.91 | 9.29 | 20 | 74.71 | 12.31 | 162.00 | 0.314 | 0.552 | -0.19 | -0.50 | 0.17 |
| HR change (last - first) | 20 | 3.13 | 5.32 | 20 | 2.92 | 4.28 | 220.00 | 0.602 | 0.657 | 0.10 | -0.26 | 0.43 |
| HRV (first 5 min) | 20 | 0.21 | 0.04 | 20 | 0.18 | 0.05 | 283.00 | 0.024 | 0.192 | 0.42 | 0.08 | 0.67 |
| HRV (last 5 min) | 20 | 0.24 | 0.11 | 20 | 0.21 | 0.06 | 212.00 | 0.758 | 0.758 | 0.06 | -0.29 | 0.40 |
| HRV change (last - first) | 20 | 0.03 | 0.10 | 20 | 0.03 | 0.08 | 169.00 | 0.414 | 0.552 | -0.16 | -0.48 | 0.20 |

**Table 4B.** Nonparametric t-tests (Mann-Whitney U) between the MindGym (Breathwork) and VR (Breathwork) groups. 'DV' refers to dependent variable. 'n' refers to group sample size. 'M' and 'SD' refer to mean and standard deviation. 'p.raw' and 'p.fdr' refer to the uncorrected p value and FDR-corrected p value. 'RBC' refers to rank biserial correlation as a measure of effect size (ranging from -1 to 1) and reflects the relative proportions of positive and negative ranks. 'RBC (low)' and 'RBC (high)' refer to the lower and upper bounds of the 95% confidence interval of RBC.

#### Table 4C (MindGym Rain vs VR Rain)

| | **MindGym (Rain)** | | | **VR (Rain)** | | | **Mann-Whitney U test** | | | | | |
|---|---|---|---|---|---|---|---|---|---|---|---|---|
| **DV** | **n** | **M** | **SD** | **n** | **M** | **SD** | **W** | **p.raw** | **p.fdr** | **RBC** | **RBC (low)** | **RBC (high)** |
| Breath Rate (first 2 min) | 18 | 16.22 | 3.07 | 16 | 15.66 | 2.93 | 159.00 | 0.616 | 0.829 | 0.10 | -0.28 | 0.46 |
| Breath Rate (last 2 min) | 18 | 16.67 | 2.18 | 16 | 17.50 | 3.46 | 110.50 | 0.254 | 0.829 | -0.23 | -0.56 | 0.16 |
| Breath Rate change (last - first) | 18 | 0.44 | 3.11 | 16 | 1.84 | 3.00 | 99.50 | 0.128 | 0.829 | -0.31 | -0.61 | 0.08 |
| EDA (first 2 min) | 25 | 41.05 | 8.62 | 22 | 38.55 | 8.44 | 315.00 | 0.403 | 0.829 | 0.15 | -0.19 | 0.45 |
| EDA (last 2 min) | 12 | 46.43 | 8.04 | 22 | 43.43 | 9.24 | 153.00 | 0.466 | 0.829 | 0.16 | -0.25 | 0.52 |
| EDA change (last - first) | 12 | 6.82 | 5.78 | 22 | 4.88 | 5.68 | 159.00 | 0.345 | 0.829 | 0.21 | -0.20 | 0.55 |
| HR (first 1 min) | 14 | 70.17 | 12.95 | 24 | 67.36 | 12.09 | 185.00 | 0.622 | 0.829 | 0.10 | -0.28 | 0.45 |
| HR (last 1 min) | 14 | 72.03 | 12.80 | 24 | 69.06 | 13.98 | 194.00 | 0.445 | 0.829 | 0.16 | -0.23 | 0.49 |
| HR change (last - first) | 14 | 1.86 | 4.26 | 24 | 1.70 | 3.07 | 177.00 | 0.800 | 0.846 | 0.05 | -0.32 | 0.41 |
| HRV (first 5 min) | 14 | 0.20 | 0.04 | 24 | 0.20 | 0.07 | 161.00 | 0.846 | 0.846 | -0.04 | -0.40 | 0.33 |
| HRV (last 5 min) | 14 | 0.20 | 0.07 | 24 | 0.19 | 0.07 | 180.00 | 0.731 | 0.846 | 0.07 | -0.30 | 0.43 |
| HRV change (last - first) | 14 | 0.01 | 0.07 | 24 | -0.01 | 0.09 | 188.00 | 0.560 | 0.829 | 0.12 | -0.26 | 0.47 |

**Table 4C.** Nonparametric t-tests (Mann-Whitney U) between the MindGym (Rain) and VR (Rain) groups. 'DV' refers to dependent variable. 'n' refers to group sample size. 'M' and 'SD' refer to mean and standard deviation. 'p.raw' and 'p.fdr' refer to the uncorrected p value and FDR-corrected p value. 'RBC' refers to rank biserial correlation as a measure of effect size (ranging from -1 to 1) and reflects the relative proportions of positive and negative ranks. 'RBC (low)' and 'RBC (high)' refer to the lower and upper bounds of the 95% confidence interval of RBC.

#### Table 4D (VR Breathwork vs VR Rain)

| | **VR (Breathwork)** | | | **VR (Rain)** | | | **Mann-Whitney U test** | | | | | |
|---|---|---|---|---|---|---|---|---|---|---|---|---|
| **DV** | **n** | **M** | **SD** | **n** | **M** | **SD** | **W** | **p.raw** | **p.fdr** | **RBC** | **RBC (low)** | **RBC (high)** |
| Breath Rate (first 2 min) | 25 | 17.74 | 3.57 | 16 | 15.66 | 2.93 | 283.00 | 0.027 | 0.108 | 0.42 | 0.08 | 0.67 |
| Breath Rate (last 2 min) | 25 | 12.06 | 3.53 | 16 | 17.50 | 3.46 | 55.00 | < .001 | < .001 | -0.73 | -0.86 | -0.50 |

| | | | | | | | | | | | |
|---|---|---|---|---|---|---|---|---|---|---|---|
| Breath Rate change (last - first) | 25 | -5.68 | 4.52 | 16 | 1.84 | 3.00 | 37.00 | < .001 | < .001 | -0.82 | -0.91 | -0.65 |
| EDA (first 2 min) | 21 | 39.51 | 12.05 | 22 | 38.55 | 8.44 | 264.00 | 0.434 | 0.473 | 0.14 | -0.20 | 0.46 |
| EDA (last 2 min) | 21 | 41.75 | 11.90 | 22 | 43.43 | 9.24 | 212.00 | 0.656 | 0.656 | -0.08 | -0.41 | 0.26 |
| EDA change (last - first) | 21 | 2.24 | 5.55 | 22 | 4.88 | 5.68 | 179.00 | 0.213 | 0.319 | -0.23 | -0.52 | 0.12 |
| HR (first 1 min) | 20 | 71.79 | 11.24 | 24 | 67.36 | 12.09 | 288.00 | 0.266 | 0.319 | 0.20 | -0.14 | 0.50 |
| HR (last 1 min) | 20 | 74.71 | 12.31 | 24 | 69.06 | 13.98 | 307.50 | 0.114 | 0.202 | 0.28 | -0.06 | 0.56 |
| HR change (last - first) | 20 | 2.92 | 4.28 | 24 | 1.70 | 3.07 | 288.00 | 0.266 | 0.319 | 0.20 | -0.14 | 0.50 |
| HRV (first 5 min) | 20 | 0.18 | 0.05 | 24 | 0.20 | 0.07 | 173.00 | 0.118 | 0.202 | -0.28 | -0.56 | 0.06 |
| HRV (last 5 min) | 20 | 0.21 | 0.06 | 24 | 0.19 | 0.07 | 310.00 | 0.102 | 0.202 | 0.29 | -0.05 | 0.57 |
| HRV change (last - first) | 20 | 0.03 | 0.08 | 24 | -0.01 | 0.09 | 325.00 | 0.046 | 0.138 | 0.35 | 0.02 | 0.62 |

**Table 4D.** Nonparametric t-tests (Mann-Whitney U) between the VR (Breathwork) and VR (Rain) groups. 'DV' refers to dependent variable. 'n' refers to group sample size. 'M' and 'SD' refer to mean and standard deviation. 'p.raw' and 'p.fdr' refer to the uncorrected p value and FDR-corrected p value. 'RBC' refers to rank biserial correlation as a measure of effect size (ranging from -1 to 1) and reflects the relative proportions of positive and negative ranks. 'RBC (low)' and 'RBC (high)' refer to the lower and upper bounds of the 95% confidence interval of RBC.

#### Table 4E (Breathwork vs Rain)

| | **Breathwork** | | | **Rain** | | | **Mann-Whitney U test** | | | | | |
|---|---|---|---|---|---|---|---|---|---|---|---|---|
| **DV** | **n** | **M** | **SD** | **n** | **M** | **SD** | **W** | **p.raw** | **p.fdr** | **RBC** | **RBC (low)** | **RBC (high)** |
| Breath Rate (first 2 min) | 52 | 18.67 | 3.98 | 34 | 15.96 | 2.97 | 1290.00 | < .001 | < .001 | 0.46 | 0.24 | 0.63 |
| Breath Rate (last 2 min) | 52 | 12.49 | 3.29 | 34 | 17.06 | 2.84 | 269.50 | < .001 | < .001 | -0.70 | -0.80 | -0.54 |
| Breath Rate change (last - first) | 52 | -6.17 | 5.08 | 34 | 1.10 | 3.09 | 198.50 | < .001 | < .001 | -0.78 | -0.86 | -0.65 |
| EDA (first 2 min) | 47 | 35.83 | 11.32 | 47 | 39.88 | 8.54 | 871.00 | 0.078 | 0.180 | -0.21 | -0.42 | 0.02 |
| EDA (last 2 min) | 36 | 40.28 | 11.84 | 34 | 44.49 | 8.83 | 491.00 | 0.158 | 0.211 | -0.20 | -0.44 | 0.07 |
| EDA change (last - first) | 36 | 3.35 | 7.11 | 34 | 5.57 | 5.71 | 488.00 | 0.148 | 0.211 | -0.20 | -0.45 | 0.07 |
| HR (first 1 min) | 40 | 69.78 | 10.90 | 38 | 68.40 | 12.31 | 808.00 | 0.637 | 0.658 | 0.06 | -0.19 | 0.31 |
| HR (last 1 min) | 40 | 72.81 | 10.93 | 38 | 70.16 | 13.46 | 871.50 | 0.267 | 0.320 | 0.15 | -0.11 | 0.39 |
| HR change (last - first) | 40 | 3.03 | 4.77 | 38 | 1.76 | 3.50 | 930.00 | 0.090 | 0.180 | 0.22 | -0.03 | 0.45 |
| HRV (first 5 min) | 40 | 0.19 | 0.04 | 38 | 0.20 | 0.06 | 715.00 | 0.658 | 0.658 | -0.06 | -0.31 | 0.20 |
| HRV (last 5 min) | 40 | 0.22 | 0.09 | 38 | 0.20 | 0.07 | 943.00 | 0.068 | 0.180 | 0.24 | -0.01 | 0.47 |
| HRV change (last - first) | 40 | 0.03 | 0.09 | 38 | 0.00 | 0.08 | 906.00 | 0.147 | 0.211 | 0.19 | -0.06 | 0.42 |

**Table 4E.** Nonparametric t-tests (Mann-Whitney U) between the Breathwork and Rain groups. 'DV' refers to dependent variable. 'n' refers to group sample size. 'M' and 'SD' refer to mean and standard deviation. 'p.raw' and 'p.fdr' refer to the uncorrected p value and FDR-corrected p value. 'RBC' refers to rank biserial correlation as a measure of effect size (ranging from -1 to 1) and reflects the relative proportions of positive and negative ranks. 'RBC (low)' and 'RBC (high)' refer to the lower and upper bounds of the 95% confidence interval of RBC.

#### Table 4F (MindGym vs VR)

| | MindGym | | | VR | | | Mann-Whitney U test | | | | | |
|---|---|---|---|---|---|---|---|---|---|---|---|---|
| DV | n | M | SD | n | M | SD | W | p.raw | p.fdr | RBC | RBC (low) | RBC (high) |
| Breath Rate (first 2 min) | 45 | 18.20 | 3.94 | 41 | 16.93 | 3.46 | 1064.50 | 0.220 | 0.660 | 0.15 | -0.09 | 0.38 |
| Breath Rate (last 2 min) | 45 | 14.40 | 3.30 | 41 | 14.18 | 4.38 | 960.50 | 0.746 | 0.956 | 0.04 | -0.20 | 0.28 |
| Breath Rate change (last - first) | 45 | -3.80 | 5.87 | 41 | -2.74 | 5.43 | 847.50 | 0.519 | 0.890 | -0.08 | -0.32 | 0.16 |
| EDA (first 2 min) | 51 | 36.88 | 10.11 | 43 | 39.02 | 10.25 | 934.00 | 0.219 | 0.660 | -0.15 | -0.37 | 0.09 |
| EDA (last 2 min) | 27 | 41.87 | 10.97 | 43 | 42.61 | 10.53 | 558.00 | 0.792 | 0.956 | -0.04 | -0.31 | 0.24 |
| EDA change (last - first) | 27 | 5.76 | 7.55 | 43 | 3.59 | 5.71 | 688.00 | 0.198 | 0.660 | 0.19 | -0.09 | 0.44 |
| HR (first 1 min) | 34 | 68.77 | 11.41 | 44 | 69.37 | 11.79 | 723.00 | 0.806 | 0.956 | -0.03 | -0.29 | 0.22 |
| HR (last 1 min) | 34 | 71.37 | 10.71 | 44 | 71.63 | 13.40 | 754.00 | 0.956 | 0.956 | 0.01 | -0.25 | 0.26 |
| HR change (last - first) | 34 | 2.60 | 4.89 | 44 | 2.26 | 3.68 | 823.00 | 0.455 | 0.890 | 0.10 | -0.16 | 0.35 |
| HRV (first 5 min) | 34 | 0.20 | 0.04 | 44 | 0.19 | 0.06 | 891.00 | 0.152 | 0.660 | 0.19 | -0.07 | 0.43 |
| HRV (last 5 min) | 34 | 0.22 | 0.09 | 44 | 0.20 | 0.07 | 819.00 | 0.480 | 0.890 | 0.10 | -0.16 | 0.34 |
| HRV change (last - first) | 34 | 0.02 | 0.09 | 44 | 0.01 | 0.09 | 732.00 | 0.877 | 0.956 | -0.02 | -0.27 | 0.23 |

**Table 4F.** Nonparametric t-tests (Mann-Whitney U) between the MindGym and VR groups. ‘DV’ refers to dependent variable. ‘n’ refers to group sample size. ‘M’ and ‘SD’ refer to mean and standard deviation. ‘p.raw’ and ‘p.fdr’ refer to the uncorrected p value and FDR-corrected p value. ‘RBC’ refers to rank biserial correlation as a measure of effect size (ranging from -1 to 1) and reflects the relative proportions of positive and negative ranks. ‘RBC (low)’ and ‘RBC (high)’ refer to the lower and upper bounds of the 95% confidence interval of RBC.
