## Supplementary material for "Contrasting cognitive, behavioral, and physiological responses to breathwork vs. naturalistic stimuli in reflective chamber and VR headset environments": S5 Tables A-J.pdf

### S5: Moderation Tables

##### Table 5A (Architex Total Score)

| DV | Tech x Stim x Moderator | | | Tech x Moderator | | | Stim x Moderator | | | Moderator | | | | |
|---|---|---|---|---|---|---|---|---|---|---|---|---|---|---|
| Architex (Total Score) | **F** | **p.raw** | **p.fdr** | **F** | **p.raw** | **p.fdr** | **F** | **p.raw** | **p.fdr** | **b** | **SE** | **Z** | **p.raw** | **p.fdr** |
| **Psych Moderator** | | | | | | | | | | | | | | |
| DPES | 0.28 | 0.598 | 0.941 | 0.98 | 0.322 | 0.661 | 5.68 | 0.017 | 0.128 | 0.15 | 0.28 | 0.55 | 0.581 | 0.846 |
| Immersion | 5.71 | 0.017 | 0.254 | 1.10 | 0.295 | 0.661 | 0.07 | 0.795 | 0.951 | -0.01 | 0.02 | -0.45 | 0.652 | 0.846 |
| IPQ (General) | 2.09 | 0.148 | 0.452 | 0.00 | 0.996 | 0.996 | 0.37 | 0.543 | 0.905 | 0.04 | 0.14 | 0.30 | 0.761 | 0.846 |
| IPQ (Involvement) | 3.29 | 0.070 | 0.380 | 0.47 | 0.492 | 0.671 | 0.16 | 0.690 | 0.941 | -0.01 | 0.06 | -0.20 | 0.846 | 0.846 |
| IPQ (Spatial Presence) | 0.01 | 0.908 | 0.941 | 2.34 | 0.126 | 0.629 | 0.02 | 0.903 | 0.968 | 0.02 | 0.07 | 0.24 | 0.808 | 0.846 |
| IPQ (Experienced Realism) | 1.37 | 0.242 | 0.560 | 0.04 | 0.834 | 0.958 | 0.44 | 0.509 | 0.905 | 0.01 | 0.07 | 0.20 | 0.844 | 0.846 |
| MODTAS | 0.09 | 0.764 | 0.941 | 0.66 | 0.418 | 0.661 | 1.50 | 0.221 | 0.827 | 0.00 | 0.01 | 0.38 | 0.708 | 0.846 |
| Openness | 2.06 | 0.151 | 0.452 | 0.02 | 0.894 | 0.958 | 5.74 | 0.017 | 0.128 | -0.01 | 0.07 | -0.20 | 0.839 | 0.846 |
| **Physio Moderator** | | | | | | | | | | | | | | |
| Motion (overall) | 0.22 | 0.639 | 0.941 | 3.55 | 0.059 | 0.446 | 0.00 | 0.968 | 0.968 | 0.04 | 0.04 | 1.03 | 0.305 | 0.819 |
| Breath Rate (first 2 min) | 0.03 | 0.860 | 0.941 | 0.72 | 0.396 | 0.661 | 0.19 | 0.662 | 0.941 | -0.08 | 0.07 | -1.09 | 0.276 | 0.819 |
| HR (first 1 min) | 3.15 | 0.076 | 0.380 | 0.59 | 0.441 | 0.661 | 4.67 | 0.031 | 0.154 | -0.02 | 0.02 | -0.98 | 0.326 | 0.819 |
| HRV (first 5 min) | 1.26 | 0.261 | 0.560 | 1.08 | 0.298 | 0.661 | 0.45 | 0.502 | 0.905 | 4.36 | 4.46 | 0.98 | 0.327 | 0.819 |
| Breath Rate change (last - first) | 0.01 | 0.941 | 0.941 | 0.22 | 0.642 | 0.803 | 0.60 | 0.439 | 0.905 | 0.06 | 0.06 | 1.11 | 0.266 | 0.819 |
| HR change (last - first) | 0.68 | 0.409 | 0.767 | 10.08 | 0.002 | 0.023 | 0.05 | 0.824 | 0.951 | -0.12 | 0.06 | -1.95 | 0.051 | 0.762 |
| HRV change (last - first) | 0.04 | 0.852 | 0.941 | 0.78 | 0.379 | 0.661 | 0.55 | 0.457 | 0.905 | -0.95 | 3.62 | -0.26 | 0.793 | 0.846 |

**Table 5A.** Results from GLM tests of moderation. 'DV' refers to dependent variable of Architex (Total Score) calculated as a difference score (post – pre). 'Psych Moderator' and 'Physio Moderator' refer to either the psychological or physiological variable defined as the moderator. 'Tech x Stim x Moderator' refers to the model of a three-way interaction between the moderator and the group factors of Tech (MindGym vs VR) and Stimuli (Breathwork vs Rain). 'Tech x Moderator' refers to the model of a two-way interaction between the moderator and the group factor of Tech (MindGym vs VR). 'Stim x Moderator' refers to the model of a two-way interaction between the moderator and the group factor of Stimuli (Breathwork vs Rain). 'Moderator' refers to the effect of the moderator from the model of only main effects. 'F' refers to F-test from the GLM model. 'b' and 'SE' refer to the unstandardized beta coefficient and its standard error. 'Z' refers to the Z-test from the model. 'p.raw' and 'p.fdr' refer to the uncorrected p value and FDR-corrected p value.

##### Table 5B (Architex Total Speed)

| DV | Tech x Stim x Moderator | | | Tech x Moderator | | | Stim x Moderator | | | Moderator | | | | |
|---|---|---|---|---|---|---|---|---|---|---|---|---|---|---|
| Architex (Total Speed) | **F** | **p.raw** | **p.fdr** | **F** | **p.raw** | **p.fdr** | **F** | **p.raw** | **p.fdr** | **b** | **SE** | **Z** | **p.raw** | **p.fdr** |
| **Psych Moderator** | | | | | | | | | | | | | | |
| DPES | 0.70 | 0.403 | 0.792 | 0.50 | 0.481 | 0.656 | 0.08 | 0.776 | 0.984 | -8.68 | 6.53 | -1.33 | 0.184 | 0.824 |
| Immersion | 0.02 | 0.887 | 0.887 | 3.05 | 0.081 | 0.394 | 0.02 | 0.881 | 0.984 | 0.54 | 0.49 | 1.09 | 0.274 | 0.824 |
| IPQ (General) | 0.18 | 0.669 | 0.837 | 1.32 | 0.251 | 0.470 | 0.18 | 0.669 | 0.984 | 2.67 | 3.46 | 0.77 | 0.440 | 0.824 |
| IPQ (Involvement) | 0.09 | 0.764 | 0.881 | 1.77 | 0.184 | 0.394 | 0.54 | 0.463 | 0.984 | 2.10 | 1.57 | 1.33 | 0.183 | 0.824 |
| IPQ (Spatial Presence) | 0.03 | 0.854 | 0.887 | 0.09 | 0.761 | 0.902 | 0.09 | 0.760 | 0.984 | -1.15 | 1.87 | -0.61 | 0.540 | 0.843 |
| IPQ (Experienced Realism) | 0.19 | 0.664 | 0.837 | 2.11 | 0.147 | 0.394 | 0.01 | 0.946 | 0.984 | 0.95 | 1.63 | 0.58 | 0.562 | 0.843 |
| MODTAS | 0.69 | 0.407 | 0.792 | 2.39 | 0.122 | 0.394 | 0.02 | 0.893 | 0.984 | -0.02 | 0.22 | -0.08 | 0.933 | 0.980 |

| | | | | | | | | | | | | | |
|---|---|---|---|---|---|---|---|---|---|---|---|---|---|
| Openness | 2.21 | 0.137 | 0.792 | 0.08 | 0.782 | 0.902 | 0.00 | 0.984 | 0.984 | -1.48 | 1.39 | -1.06 | 0.289 | 0.824 |
| **Physio Moderator** | | | | | | | | | | | | | | |
| Motion (overall) | 0.74 | 0.390 | 0.792 | 1.91 | 0.167 | 0.394 | 6.83 | 0.009 | 0.068 | -0.10 | 1.01 | -0.10 | 0.924 | 0.980 |
| Breath Rate (first 2 min) | 0.94 | 0.332 | 0.792 | 0.65 | 0.419 | 0.628 | 0.53 | 0.465 | 0.984 | 0.55 | 1.86 | 0.30 | 0.767 | 0.980 |
| HR (first 1 min) | 0.34 | 0.559 | 0.837 | 1.87 | 0.171 | 0.394 | 1.03 | 0.311 | 0.984 | 0.55 | 0.66 | 0.84 | 0.403 | 0.824 |
| HRV (first 5 min) | 2.07 | 0.156 | 0.792 | 6.50 | 0.013 | 0.195 | 8.49 | 0.005 | 0.068 | 344.90 | 132.69 | 2.60 | 0.009 | 0.140 |
| Breath Rate change (last - first) | 0.41 | 0.522 | 0.837 | 0.01 | 0.906 | 0.906 | 0.96 | 0.328 | 0.984 | 1.41 | 1.56 | 0.91 | 0.365 | 0.824 |
| HR change (last - first) | 0.64 | 0.423 | 0.792 | 0.04 | 0.848 | 0.906 | 0.03 | 0.863 | 0.984 | 0.04 | 1.57 | 0.03 | 0.980 | 0.980 |
| HRV change (last - first) | 65.00 | 0.300 | 0.792 | 68.00 | 0.386 | 0.628 | 68.00 | 0.676 | 0.984 | 9.84 | 82.18 | 0.12 | 0.905 | 0.980 |

**Table 5B.** Results from GLM tests of moderation. 'DV' refers to dependent variable of Architex (Total Speed) calculated as a difference score (post – pre). 'Psych Moderator' and 'Physio Moderator' refer to either the psychological or physiological variable defined as the moderator. 'Tech x Stim x Moderator' refers to the model of a three-way interaction between the moderator and the group factors of Tech (MindGym vs VR) and Stimuli (Breathwork vs Rain). 'Tech x Moderator' refers to the model of a two-way interaction between the moderator and the group factor of Tech (MindGym vs VR). 'Stim x Moderator' refers to the model of a two-way interaction between the moderator and the group factor of Stimuli (Breathwork vs Rain). 'Moderator' refers to the effect of the moderator from the model of only main effects. 'F' refers to F-test from the GLM model. 'b' and 'SE' refer to the unstandardized beta coefficient and its standard error. 'Z' refers to the Z-test from the model. 'p.raw' and 'p.fdr' refer to the uncorrected p value and FDR-corrected p value.

#### Table 5C (TMT RTACC)

| **DV** | **Tech x Stim x Moderator** | | | **Tech x Moderator** | | | **Stim x Moderator** | | | **Moderator** | | | | |
|---|---|---|---|---|---|---|---|---|---|---|---|---|---|---|
| TMT (RTACC) | **F** | **p.raw** | **p.fdr** | **F** | **p.raw** | **p.fdr** | **F** | **p.raw** | **p.fdr** | **b** | **SE** | **Z** | **p.raw** | **p.fdr** |
| **Psych Moderator** | | | | | | | | | | | | | | |
| DPES | 0.72 | 0.397 | 0.725 | 0.27 | 0.601 | 0.772 | 0.31 | 0.581 | 0.686 | -7.74 | 4.33 | -1.79 | 0.074 | 0.335 |
| Immersion | 0.04 | 0.850 | 0.850 | 0.30 | 0.582 | 0.772 | 4.17 | 0.041 | 0.529 | 0.25 | 0.27 | 0.92 | 0.357 | 0.669 |
| IPQ (General) | 1.84 | 0.175 | 0.609 | 0.02 | 0.895 | 0.895 | 0.54 | 0.463 | 0.686 | 1.77 | 2.16 | 0.82 | 0.414 | 0.689 |
| IPQ (Involvement) | 6.66 | 0.010 | 0.149 | 0.22 | 0.642 | 0.772 | 0.28 | 0.594 | 0.686 | 1.23 | 0.87 | 1.41 | 0.157 | 0.473 |
| IPQ (Spatial Presence) | 0.19 | 0.664 | 0.850 | 0.46 | 0.496 | 0.772 | 0.53 | 0.465 | 0.686 | 1.40 | 1.22 | 1.15 | 0.252 | 0.612 |
| IPQ (Experienced Realism) | 1.22 | 0.270 | 0.609 | 2.63 | 0.105 | 0.759 | 1.39 | 0.238 | 0.595 | 2.08 | 1.06 | 1.96 | 0.050 | 0.335 |
| MODTAS | 0.24 | 0.624 | 0.850 | 0.77 | 0.380 | 0.772 | 0.33 | 0.564 | 0.686 | -0.23 | 0.13 | -1.70 | 0.089 | 0.335 |
| Openness | 0.61 | 0.435 | 0.725 | 0.18 | 0.673 | 0.772 | 0.00 | 0.998 | 0.998 | 0.15 | 0.99 | 0.15 | 0.878 | 0.940 |
| **Physio Moderator** | | | | | | | | | | | | | | |
| Motion (overall) | 0.06 | 0.811 | 0.850 | 1.63 | 0.202 | 0.759 | 2.74 | 0.098 | 0.529 | -0.31 | 0.49 | -0.63 | 0.528 | 0.720 |
| Breath Rate (first 2 min) | 1.15 | 0.284 | 0.609 | 0.13 | 0.720 | 0.772 | 1.25 | 0.264 | 0.595 | 0.93 | 0.87 | 1.07 | 0.286 | 0.612 |
| HR (first 1 min) | 0.12 | 0.728 | 0.850 | 0.13 | 0.720 | 0.772 | 1.26 | 0.262 | 0.595 | 0.12 | 0.30 | 0.39 | 0.695 | 0.869 |
| HRV (first 5 min) | 3.41 | 0.065 | 0.324 | 3.62 | 0.057 | 0.759 | 0.44 | 0.505 | 0.686 | 131.00 | 53.94 | 2.43 | 0.015 | 0.227 |
| Breath Rate change (last - first) | 1.35 | 0.245 | 0.609 | 0.86 | 0.354 | 0.772 | 2.62 | 0.106 | 0.529 | -0.04 | 0.74 | -0.05 | 0.957 | 0.957 |
| HR change (last - first) | 0.08 | 0.776 | 0.850 | 0.53 | 0.468 | 0.772 | 1.18 | 0.278 | 0.595 | 0.67 | 0.94 | 0.72 | 0.475 | 0.712 |
| HRV change (last - first) | 63.00 | 0.038 | 0.287 | 66.00 | 0.167 | 0.759 | 0.00 | 0.961 | 0.998 | -11.11 | 39.32 | -0.28 | 0.777 | 0.897 |

**Table 5C.** Results from GLM tests of moderation. 'DV' refers to dependent variable of TMT (RTACC) calculated as a difference score (post – pre). 'Psych Moderator' and 'Physio Moderator' refer to either the psychological or physiological variable defined as the moderator. 'Tech x Stim x Moderator' refers to the model of a three-way interaction between the moderator and the group factors of Tech (MindGym vs VR) and Stimuli (Breathwork vs Rain). 'Tech x Moderator' refers to the model of a two-way interaction between the moderator and the group factor of Tech (MindGym vs VR). 'Stim x Moderator' refers to the model of a two-way interaction between the moderator and the group factor of Stimuli (Breathwork vs Rain). 'Moderator' refers to the effect of the moderator from the model of only main effects. ‘F’ refers to F-test from the GLM model. ‘b’ and ‘SE’ refer to the unstandardized beta coefficient and its standard error. ‘Z’ refers to the Z-test from the model. ‘p.raw’ and ‘p.fdr’ refer to the uncorrected p value and FDR-corrected p value.

#### Table 5D (Arousal)

| DV | Tech x Stim x Moderator | | | Tech x Moderator | | | Stim x Moderator | | | Moderator | | | | |
|---|---|---|---|---|---|---|---|---|---|---|---|---|---|---|
| Arousal | F | p.raw | p.fdr | F | p.raw | p.fdr | F | p.raw | p.fdr | b | SE | Z | p.raw | p.fdr |
| **Psych Moderator** | | | | | | | | | | | | | | |
| DPES | 0.42 | 0.518 | 0.948 | 2.29 | 0.130 | 0.456 | 0.00 | 0.983 | 0.983 | 0.01 | 0.25 | 0.04 | 0.970 | 0.980 |
| Immersion | 0.56 | 0.454 | 0.948 | 0.00 | 0.982 | 0.982 | 1.16 | 0.281 | 0.919 | 0.00 | 0.02 | 0.04 | 0.969 | 0.980 |
| IPQ (General) | 0.03 | 0.875 | 0.948 | 1.42 | 0.233 | 0.545 | 0.63 | 0.428 | 0.983 | 0.11 | 0.13 | 0.79 | 0.428 | 0.980 |
| IPQ (Involvement) | 0.82 | 0.366 | 0.948 | 2.05 | 0.152 | 0.456 | 0.02 | 0.877 | 0.983 | 0.01 | 0.06 | 0.16 | 0.870 | 0.980 |
| IPQ (Spatial Presence) | 0.05 | 0.826 | 0.948 | 0.16 | 0.685 | 0.857 | 3.35 | 0.067 | 0.918 | -0.07 | 0.07 | -1.04 | 0.298 | 0.980 |
| IPQ (Experienced Realism) | 1.70 | 0.192 | 0.720 | 1.13 | 0.287 | 0.545 | 0.01 | 0.940 | 0.983 | 0.00 | 0.06 | 0.05 | 0.960 | 0.980 |
| MODTAS | 0.63 | 0.426 | 0.948 | 2.14 | 0.144 | 0.456 | 0.03 | 0.854 | 0.983 | -0.01 | 0.01 | -0.82 | 0.415 | 0.980 |
| Openness | 1.72 | 0.190 | 0.720 | 2.38 | 0.123 | 0.456 | 1.29 | 0.256 | 0.919 | -0.03 | 0.05 | -0.52 | 0.606 | 0.980 |
| **Physio Moderator** | | | | | | | | | | | | | | |
| Motion (overall) | 0.01 | 0.934 | 0.948 | 1.12 | 0.291 | 0.545 | 0.01 | 0.923 | 0.983 | -0.01 | 0.03 | -0.34 | 0.731 | 0.980 |
| Breath Rate (first 2 min) | 0.21 | 0.649 | 0.948 | 0.22 | 0.641 | 0.857 | 0.16 | 0.694 | 0.983 | 0.00 | 0.06 | -0.03 | 0.980 | 0.980 |
| HR (first 1 min) | 0.00 | 0.948 | 0.948 | 0.11 | 0.744 | 0.859 | 0.00 | 0.961 | 0.983 | 0.03 | 0.03 | 1.26 | 0.209 | 0.980 |
| HRV (first 5 min) | 0.07 | 0.792 | 0.948 | 0.64 | 0.426 | 0.709 | 1.05 | 0.306 | 0.919 | -2.18 | 5.56 | -0.39 | 0.696 | 0.980 |
| Breath Rate change (last - first) | 2.61 | 0.106 | 0.720 | 0.01 | 0.935 | 0.982 | 0.05 | 0.825 | 0.983 | 0.02 | 0.05 | 0.43 | 0.670 | 0.980 |
| HR change (last - first) | 66.00 | 0.845 | 0.948 | 69.00 | 0.655 | 0.857 | 69.00 | 0.122 | 0.918 | 0.08 | 0.07 | 1.22 | 0.221 | 0.980 |
| HRV change (last - first) | 2.27 | 0.132 | 0.720 | 6.74 | 0.009 | 0.141 | 0.31 | 0.578 | 0.983 | 2.09 | 3.51 | 0.60 | 0.551 | 0.980 |

**Table 5D.** Results from GLM tests of moderation. ‘DV’ refers to dependent variable of Arousal calculated as a difference score (post – pre). ‘Psych Moderator’ and ‘Physio Moderator’ refer to either the psychological or physiological variable defined as the moderator. ‘Tech x Stim x Moderator’ refers to the model of a three-way interaction between the moderator and the group factors of Tech (MindGym vs VR) and Stimuli (Breathwork vs Rain). ‘Tech x Moderator’ refers to the model of a two-way interaction between the moderator and the group factor of Tech (MindGym vs VR). ‘Stim x Moderator’ refers to the model of a two-way interaction between the moderator and the group factor of Stimuli (Breathwork vs Rain). ‘Moderator’ refers to the effect of the moderator from the model of only main effects. ‘F’ refers to F-test from the GLM model. ‘b’ and ‘SE’ refer to the unstandardized beta coefficient and its standard error. ‘Z’ refers to the Z-test from the model. ‘p.raw’ and ‘p.fdr’ refer to the uncorrected p value and FDR-corrected p value.

#### Table 5E (POMS)

| DV | Tech x Stim x Moderator | | | Tech x Moderator | | | Stim x Moderator | | | Moderator | | | | |
|---|---|---|---|---|---|---|---|---|---|---|---|---|---|---|
| POMS | F | p.raw | p.fdr | F | p.raw | p.fdr | F | p.raw | p.fdr | b | SE | Z | p.raw | p.fdr |
| **Psych Moderator** | | | | | | | | | | | | | | |
| DPES | 0.05 | 0.831 | 0.891 | 0.74 | 0.390 | 0.824 | 1.62 | 0.206 | 0.806 | -0.86 | 1.65 | -0.52 | 0.603 | 0.886 |
| Immersion | 0.07 | 0.795 | 0.891 | 0.53 | 0.466 | 0.824 | 0.01 | 0.923 | 0.940 | -0.04 | 0.11 | -0.35 | 0.728 | 0.911 |
| IPQ (General) | 0.37 | 0.543 | 0.757 | 2.67 | 0.105 | 0.674 | 0.33 | 0.568 | 0.806 | 0.81 | 0.80 | 1.01 | 0.313 | 0.886 |
| IPQ (Involvement) | 3.05 | 0.084 | 0.599 | 0.95 | 0.331 | 0.824 | 0.29 | 0.591 | 0.806 | -0.04 | 0.35 | -0.10 | 0.918 | 0.918 |
| IPQ (Spatial Presence) | 2.32 | 0.131 | 0.599 | 0.59 | 0.445 | 0.824 | 0.33 | 0.569 | 0.806 | -0.24 | 0.43 | -0.55 | 0.581 | 0.886 |
| IPQ (Experienced Realism) | 0.55 | 0.462 | 0.757 | 0.01 | 0.933 | 0.933 | 0.02 | 0.892 | 0.940 | -0.18 | 0.39 | -0.46 | 0.649 | 0.886 |
| MODTAS | 0.80 | 0.374 | 0.701 | 2.35 | 0.128 | 0.674 | 0.21 | 0.645 | 0.806 | -0.01 | 0.05 | -0.16 | 0.871 | 0.918 |

| | | | | | | | | | | | | | | |
|---|---|---|---|---|---|---|---|---|---|---|---|---|---|---|
| Openness | 0.80 | 0.373 | 0.701 | 0.24 | 0.628 | 0.824 | 0.30 | 0.588 | 0.806 | -0.22 | 0.37 | -0.60 | 0.551 | 0.886 |
| **Physio Moderator** | | | | | | | | | | | | | | |
| Motion (overall) | 0.06 | 0.808 | 0.891 | 0.13 | 0.721 | 0.824 | 1.05 | 0.307 | 0.806 | 0.42 | 0.22 | 1.93 | 0.053 | 0.398 |
| Breath Rate (first 2 min) | 1.26 | 0.265 | 0.662 | 0.18 | 0.669 | 0.824 | 1.28 | 0.261 | 0.806 | -0.42 | 0.41 | -1.01 | 0.311 | 0.886 |
| HR (first 1 min) | 2.23 | 0.140 | 0.599 | 0.12 | 0.732 | 0.824 | 0.28 | 0.597 | 0.806 | -0.03 | 0.15 | -0.18 | 0.855 | 0.918 |
| HRV (first 5 min) | 0.01 | 0.934 | 0.934 | 0.82 | 0.368 | 0.824 | 0.01 | 0.940 | 0.940 | 21.22 | 33.79 | 0.63 | 0.530 | 0.886 |
| Breath Rate change (last - first) | 1.68 | 0.200 | 0.599 | 0.19 | 0.661 | 0.824 | 0.30 | 0.584 | 0.806 | 0.45 | 0.32 | 1.38 | 0.169 | 0.843 |
| HR change (last - first) | 1.77 | 0.184 | 0.599 | 2.24 | 0.135 | 0.674 | 2.32 | 0.128 | 0.806 | -0.22 | 0.41 | -0.52 | 0.601 | 0.886 |
| HRV change (last - first) | 0.35 | 0.555 | 0.757 | 0.09 | 0.769 | 0.824 | 0.74 | 0.391 | 0.806 | -38.21 | 18.29 | -2.09 | 0.037 | 0.398 |

**Table 5E.** Results from GLM tests of moderation. 'DV' refers to dependent variable of POMS (Total mood disturbance) calculated as a difference score (post – pre). 'Psych Moderator' and 'Physio Moderator' refer to either the psychological or physiological variable defined as the moderator. 'Tech x Stim x Moderator' refers to the model of a three-way interaction between the moderator and the group factors of Tech (MindGym vs VR) and Stimuli (Breathwork vs Rain). 'Tech x Moderator' refers to the model of a two-way interaction between the moderator and the group factor of Tech (MindGym vs VR). 'Stim x Moderator' refers to the model of a two-way interaction between the moderator and the group factor of Stimuli (Breathwork vs Rain). 'Moderator' refers to the effect of the moderator from the model of only main effects. 'F' refers to F-test from the GLM model. 'b' and 'SE' refer to the unstandardized beta coefficient and its standard error. 'Z' refers to the Z-test from the model. 'p.raw' and 'p.fdr' refer to the uncorrected p value and FDR-corrected p value.

#### Table 5F (STAI)

| **DV** | **Tech x Stim x Moderator** | | | **Tech x Moderator** | | | **Stim x Moderator** | | | **Moderator** | | | | |
|---|---|---|---|---|---|---|---|---|---|---|---|---|---|---|
| Anxiety (STAI) | **F** | **p.raw** | **p.fdr** | **F** | **p.raw** | **p.fdr** | **F** | **p.raw** | **p.fdr** | **b** | **SE** | **Z** | **p.raw** | **p.fdr** |
| **Psych Moderator** | | | | | | | | | | | | | | |
| DPES | 0.04 | 0.842 | 0.985 | 1.98 | 0.162 | 0.608 | 2.71 | 0.103 | 0.411 | -0.35 | 0.76 | -0.46 | 0.647 | 0.814 |
| Immersion | 0.53 | 0.467 | 0.985 | 0.37 | 0.545 | 0.817 | 0.99 | 0.322 | 0.411 | -0.05 | 0.05 | -1.01 | 0.313 | 0.740 |
| IPQ (General) | 0.00 | 0.963 | 0.985 | 2.00 | 0.160 | 0.608 | 1.20 | 0.275 | 0.411 | -0.42 | 0.37 | -1.16 | 0.247 | 0.740 |
| IPQ (Involvement) | 0.03 | 0.858 | 0.985 | 0.12 | 0.729 | 0.831 | 0.17 | 0.681 | 0.681 | 0.03 | 0.16 | 0.17 | 0.867 | 0.929 |
| IPQ (Spatial Presence) | 0.34 | 0.564 | 0.985 | 2.37 | 0.127 | 0.608 | 0.25 | 0.621 | 0.681 | -0.16 | 0.19 | -0.83 | 0.407 | 0.740 |
| IPQ (Experienced Realism) | 0.02 | 0.876 | 0.985 | 1.09 | 0.299 | 0.699 | 1.15 | 0.285 | 0.411 | -0.12 | 0.18 | -0.69 | 0.493 | 0.740 |
| MODTAS | 0.10 | 0.758 | 0.985 | 0.62 | 0.432 | 0.720 | 6.42 | 0.013 | 0.095 | -0.04 | 0.02 | -1.67 | 0.095 | 0.357 |
| Openness | 0.58 | 0.450 | 0.985 | 0.97 | 0.327 | 0.699 | 9.58 | 0.003 | 0.038 | -0.27 | 0.16 | -1.68 | 0.093 | 0.357 |
| **Physio Moderator** | | | | | | | | | | | | | | |
| Motion (overall) | 1.45 | 0.232 | 0.985 | 5.72 | 0.018 | 0.276 | 1.43 | 0.235 | 0.411 | 0.07 | 0.10 | 0.72 | 0.470 | 0.740 |
| Breath Rate (first 2 min) | 0.05 | 0.817 | 0.985 | 0.08 | 0.785 | 0.831 | 1.81 | 0.182 | 0.411 | -0.56 | 0.19 | -2.90 | 0.004 | 0.057 |
| HR (first 1 min) | 0.01 | 0.915 | 0.985 | 0.12 | 0.732 | 0.831 | 0.23 | 0.636 | 0.681 | 0.00 | 0.08 | -0.04 | 0.969 | 0.969 |
| HRV (first 5 min) | 0.00 | 0.985 | 0.985 | 0.05 | 0.831 | 0.831 | 0.97 | 0.329 | 0.411 | -6.19 | 16.37 | -0.38 | 0.705 | 0.814 |
| Breath Rate change (last - first) | 2.20 | 0.143 | 0.985 | 0.21 | 0.646 | 0.831 | 2.40 | 0.126 | 0.411 | 0.39 | 0.15 | 2.50 | 0.013 | 0.094 |
| HR change (last - first) | 0.51 | 0.479 | 0.985 | 0.81 | 0.373 | 0.699 | 1.88 | 0.175 | 0.411 | -0.15 | 0.20 | -0.76 | 0.446 | 0.740 |
| HRV change (last - first) | 0.06 | 0.815 | 0.985 | 1.33 | 0.253 | 0.699 | 1.25 | 0.268 | 0.411 | 3.98 | 9.53 | 0.42 | 0.677 | 0.814 |

**Table 5F.** Results from GLM tests of moderation. 'DV' refers to dependent variable of STAI calculated as a difference score (post – pre). 'Psych Moderator' and 'Physio Moderator' refer to either the psychological or physiological variable defined as the moderator. 'Tech x Stim x Moderator' refers to the model of a three-way interaction between the moderator and the group factors of Tech (MindGym vs VR) and Stimuli (Breathwork vs Rain). 'Tech x Moderator' refers to the model of a two-way interaction between the moderator and the group factor of Tech (MindGym vs VR). 'Stim x Moderator' refers to the model of a two-way interaction between the moderator and the group factor of Stimuli (Breathwork vs Rain). 'Moderator' refers to the effect of the moderator from the model of only main effects. 'F' refers to F-test from the GLM model. 'b' and 'SE' refer to the unstandardized beta coefficient and its standard error. 'Z' refers to the Z-test from the model. 'p.raw' and 'p.fdr' refer to the uncorrected p value and FDR-corrected p value.

#### Table 5G (Valence)

| DV | Tech x Stim x Moderator | | | Tech x Moderator | | | Stim x Moderator | | | Moderator | | | | |
|---|---|---|---|---|---|---|---|---|---|---|---|---|---|---|
| Valence | **F** | **p.raw** | **p.fdr** | **F** | **p.raw** | **p.fdr** | **F** | **p.raw** | **p.fdr** | **b** | **SE** | **Z** | **p.raw** | **p.fdr** |
| **Psych Moderator** | | | | | | | | | | | | | | |
| DPES | 0.02 | 0.894 | 0.915 | 0.34 | 0.563 | 0.876 | 0.12 | 0.733 | 0.909 | -0.07 | 0.20 | -0.33 | 0.742 | 0.823 |
| Immersion | 1.48 | 0.227 | 0.710 | 0.10 | 0.759 | 0.876 | 0.01 | 0.909 | 0.909 | -0.01 | 0.01 | -0.99 | 0.323 | 0.813 |
| IPQ (General) | 0.09 | 0.769 | 0.915 | 2.50 | 0.117 | 0.876 | 0.47 | 0.496 | 0.909 | -0.05 | 0.10 | -0.45 | 0.651 | 0.813 |
| IPQ (Involvement) | 3.46 | 0.066 | 0.583 | 1.82 | 0.181 | 0.876 | 0.08 | 0.775 | 0.909 | 0.00 | 0.04 | 0.01 | 0.995 | 0.995 |
| IPQ (Spatial Presence) | 0.01 | 0.915 | 0.915 | 0.10 | 0.748 | 0.876 | 0.69 | 0.407 | 0.909 | 0.03 | 0.05 | 0.50 | 0.619 | 0.813 |
| IPQ (Experienced Realism) | 0.21 | 0.652 | 0.915 | 0.19 | 0.663 | 0.876 | 0.46 | 0.501 | 0.909 | 0.03 | 0.05 | 0.63 | 0.526 | 0.813 |
| MODTAS | 0.10 | 0.758 | 0.915 | 1.48 | 0.226 | 0.876 | 1.17 | 0.283 | 0.909 | 0.00 | 0.01 | -0.30 | 0.768 | 0.823 |
| Openness | 0.32 | 0.574 | 0.915 | 0.56 | 0.458 | 0.876 | 0.76 | 0.385 | 0.909 | 0.05 | 0.05 | 1.04 | 0.299 | 0.813 |
| **Physio Moderator** | | | | | | | | | | | | | | |
| Motion (overall) | 0.65 | 0.421 | 0.807 | 1.38 | 0.244 | 0.876 | 1.34 | 0.249 | 0.909 | -0.05 | 0.02 | -1.93 | 0.054 | 0.402 |
| Breath Rate (first 2 min) | 0.63 | 0.431 | 0.807 | 0.00 | 0.987 | 0.987 | 0.90 | 0.346 | 0.909 | -0.03 | 0.05 | -0.51 | 0.607 | 0.813 |
| HR (first 1 min) | 0.02 | 0.903 | 0.915 | 0.11 | 0.743 | 0.876 | 0.30 | 0.588 | 0.909 | 0.01 | 0.02 | 0.51 | 0.611 | 0.813 |
| HRV (first 5 min) | 3.22 | 0.078 | 0.583 | 0.01 | 0.937 | 0.987 | 0.04 | 0.836 | 0.909 | 11.13 | 4.24 | 2.62 | 0.009 | 0.131 |
| Breath Rate change (last - first) | 1.54 | 0.220 | 0.710 | 0.77 | 0.384 | 0.876 | 0.13 | 0.715 | 0.909 | -0.02 | 0.04 | -0.60 | 0.551 | 0.813 |
| HR change (last - first) | 0.91 | 0.343 | 0.807 | 1.11 | 0.296 | 0.876 | 0.02 | 0.898 | 0.909 | 0.03 | 0.04 | 0.62 | 0.536 | 0.813 |
| HRV change (last - first) | 1.43 | 0.237 | 0.710 | 0.13 | 0.725 | 0.876 | 0.10 | 0.756 | 0.909 | -3.21 | 2.47 | -1.30 | 0.194 | 0.813 |

**Table 5G.** Results from GLM tests of moderation. 'DV' refers to dependent variable of Valence calculated as a difference score (post – pre). 'Psych Moderator' and 'Physio Moderator' refer to either the psychological or physiological variable defined as the moderator. 'Tech x Stim x Moderator' refers to the model of a three-way interaction between the moderator and the group factors of Tech (MindGym vs VR) and Stimuli (Breathwork vs Rain). 'Tech x Moderator' refers to the model of a two-way interaction between the moderator and the group factor of Tech (MindGym vs VR). 'Stim x Moderator' refers to the model of a two-way interaction between the moderator and the group factor of Stimuli (Breathwork vs Rain). 'Moderator' refers to the effect of the moderator from the model of only main effects. 'F' refers to F-test from the GLM model. 'b' and 'SE' refer to the unstandardized beta coefficient and its standard error. 'Z' refers to the Z-test from the model. 'p.raw' and 'p.fdr' refer to the uncorrected p value and FDR-corrected p value.

#### Table 5H (Awe)

| DV | Tech x Stim x Moderator | | | Tech x Moderator | | | Stim x Moderator | | | Moderator | | | | |
|---|---|---|---|---|---|---|---|---|---|---|---|---|---|---|
| Awe | **F** | **p.raw** | **p.fdr** | **F** | **p.raw** | **p.fdr** | **F** | **p.raw** | **p.fdr** | **b** | **SE** | **Z** | **p.raw** | **p.fdr** |
| **Psych Moderator** | | | | | | | | | | | | | | |
| DPES | 1.74 | 0.190 | 0.824 | 0.14 | 0.711 | 0.887 | 0.20 | 0.657 | 0.957 | 13.89 | 3.48 | 4.00 | < .0001 | < .001 |
| Immersion | 0.03 | 0.870 | 0.892 | 0.08 | 0.776 | 0.887 | 0.02 | 0.896 | 0.957 | 1.16 | 0.22 | 5.17 | < .0001 | < .001 |
| IPQ (General) | 0.02 | 0.892 | 0.892 | 4.70 | 0.032 | 0.362 | 1.92 | 0.169 | 0.853 | 6.18 | 1.72 | 3.59 | < .001 | 0.002 |
| IPQ (Involvement) | 1.08 | 0.301 | 0.892 | 0.12 | 0.727 | 0.887 | 0.82 | 0.367 | 0.853 | 2.24 | 0.76 | 2.96 | 0.003 | 0.009 |
| IPQ (Spatial Presence) | 0.38 | 0.538 | 0.892 | 0.00 | 0.957 | 0.957 | 0.77 | 0.381 | 0.853 | 1.83 | 0.93 | 1.98 | 0.048 | 0.104 |
| IPQ (Experienced Realism) | 1.52 | 0.220 | 0.824 | 0.34 | 0.560 | 0.887 | 3.98 | 0.048 | 0.725 | 0.68 | 0.87 | 0.78 | 0.436 | 0.620 |

| | | | | | | | | | | | | | | |
|---|---|---|---|---|---|---|---|---|---|---|---|---|---|---|
| MODTAS | 0.56 | 0.458 | 0.892 | 0.05 | 0.828 | 0.887 | 0.38 | 0.539 | 0.899 | 0.37 | 0.11 | 3.28 | 0.001 | 0.004 |
| Openness | 0.36 | 0.553 | 0.892 | 0.85 | 0.359 | 0.796 | 0.03 | 0.871 | 0.957 | 1.22 | 0.81 | 1.51 | 0.131 | 0.218 |
| **Physio Moderator** | | | | | | | | | | | | | | |
| Motion (overall) | 2.69 | 0.104 | 0.824 | 0.14 | 0.711 | 0.887 | 1.41 | 0.238 | 0.853 | 0.79 | 0.50 | 1.59 | 0.112 | 0.210 |
| Breath Rate (first 2 min) | 0.13 | 0.724 | 0.892 | 1.08 | 0.301 | 0.796 | 0.06 | 0.805 | 0.957 | 0.15 | 1.02 | 0.15 | 0.883 | 0.883 |
| HR (first 1 min) | 2.45 | 0.122 | 0.824 | 0.59 | 0.446 | 0.837 | 0.01 | 0.908 | 0.957 | -0.70 | 0.29 | -2.39 | 0.017 | 0.042 |
| HRV (first 5 min) | 0.03 | 0.853 | 0.892 | 4.04 | 0.048 | 0.362 | 0.00 | 0.957 | 0.957 | 42.30 | 65.70 | 0.64 | 0.520 | 0.650 |
| Breath Rate change (last - first) | 0.02 | 0.876 | 0.892 | 0.81 | 0.371 | 0.796 | 0.82 | 0.367 | 0.853 | -0.19 | 0.82 | -0.24 | 0.814 | 0.872 |
| HR change (last - first) | 0.17 | 0.686 | 0.892 | 2.44 | 0.123 | 0.508 | 0.72 | 0.398 | 0.853 | -0.60 | 0.80 | -0.75 | 0.455 | 0.620 |
| HRV change (last - first) | 0.68 | 0.412 | 0.892 | 2.28 | 0.135 | 0.508 | 0.55 | 0.461 | 0.865 | 9.92 | 38.69 | 0.26 | 0.798 | 0.872 |

**Table 5H.** Results from GLM tests of moderation. 'DV' refers to dependent variable of Awe from the post timepoint. 'Psych Moderator' and 'Physio Moderator' refer to either the psychological or physiological variable defined as the moderator. 'Tech x Stim x Moderator' refers to the model of a three-way interaction between the moderator and the group factors of Tech (MindGym vs VR) and Stimuli (Breathwork vs Rain). 'Tech x Moderator' refers to the model of a two-way interaction between the moderator and the group factor of Tech (MindGym vs VR). 'Stim x Moderator' refers to the model of a two-way interaction between the moderator and the group factor of Stimuli (Breathwork vs Rain). 'Moderator' refers to the effect of the moderator from the model of only main effects. 'F' refers to F-test from the GLM model. 'b' and 'SE' refer to the unstandardized beta coefficient and its standard error. 'Z' refers to the Z-test from the model. 'p.raw' and 'p.fdr' refer to the uncorrected p value and FDR-corrected p value.

#### Table 5I (EDI)

| **DV** | **Tech x Stim x Moderator** | | | **Tech x Moderator** | | | **Stim x Moderator** | | | **Moderator** | | | | |
|---|---|---|---|---|---|---|---|---|---|---|---|---|---|---|
| EDI | **F** | **p.raw** | **p.fdr** | **F** | **p.raw** | **p.fdr** | **F** | **p.raw** | **p.fdr** | **b** | **SE** | **Z** | **p.raw** | **p.fdr** |
| **Psych Moderator** | | | | | | | | | | | | | | |
| DPES | 0.25 | 0.617 | 0.891 | 0.55 | 0.461 | 0.894 | 1.09 | 0.299 | 0.499 | 0.54 | 0.27 | 2.03 | 0.043 | 0.128 |
| Immersion | 0.63 | 0.431 | 0.891 | 0.00 | 0.996 | 0.996 | 2.95 | 0.089 | 0.308 | 0.07 | 0.02 | 4.55 | < .0001 | < .001 |
| IPQ (General) | 0.00 | 0.949 | 0.949 | 4.05 | 0.047 | 0.351 | 5.88 | 0.017 | 0.257 | 0.32 | 0.13 | 2.49 | 0.013 | 0.048 |
| IPQ (Involvement) | 0.05 | 0.831 | 0.891 | 0.21 | 0.646 | 0.969 | 0.08 | 0.784 | 0.904 | 0.18 | 0.05 | 3.24 | 0.001 | 0.009 |
| IPQ (Spatial Presence) | 2.98 | 0.088 | 0.459 | 2.21 | 0.140 | 0.526 | 0.82 | 0.367 | 0.551 | 0.09 | 0.06 | 1.39 | 0.165 | 0.221 |
| IPQ (Experienced Realism) | 2.90 | 0.092 | 0.459 | 0.41 | 0.521 | 0.894 | 2.71 | 0.103 | 0.308 | 0.09 | 0.06 | 1.45 | 0.147 | 0.221 |
| MODTAS | 0.06 | 0.806 | 0.891 | 0.01 | 0.941 | 0.996 | 0.00 | 0.971 | 0.971 | 0.03 | 0.01 | 2.98 | 0.003 | 0.015 |
| Openness | 0.05 | 0.823 | 0.891 | 0.02 | 0.886 | 0.996 | 3.54 | 0.063 | 0.308 | 0.02 | 0.06 | 0.29 | 0.772 | 0.828 |
| **Physio Moderator** | | | | | | | | | | | | | | |
| Motion (overall) | 2.08 | 0.153 | 0.573 | 0.03 | 0.862 | 0.996 | 1.76 | 0.188 | 0.402 | -0.07 | 0.04 | -1.83 | 0.067 | 0.168 |
| Breath Rate (first 2 min) | 4.48 | 0.038 | 0.459 | 5.40 | 0.023 | 0.345 | 1.93 | 0.170 | 0.402 | 0.10 | 0.07 | 1.35 | 0.177 | 0.221 |
| HR (first 1 min) | 1.25 | 0.269 | 0.673 | 0.79 | 0.379 | 0.894 | 1.16 | 0.286 | 0.499 | -0.04 | 0.02 | -1.60 | 0.110 | 0.203 |
| HRV (first 5 min) | 0.05 | 0.821 | 0.891 | 0.40 | 0.528 | 0.894 | 0.54 | 0.465 | 0.586 | 7.86 | 4.96 | 1.59 | 0.113 | 0.203 |
| Breath Rate change (last - first) | 1.58 | 0.213 | 0.639 | 0.11 | 0.745 | 0.996 | 0.53 | 0.469 | 0.586 | -0.09 | 0.06 | -1.55 | 0.122 | 0.203 |
| HR change (last - first) | 0.13 | 0.717 | 0.891 | 2.51 | 0.119 | 0.526 | 4.04 | 0.049 | 0.308 | -0.05 | 0.06 | -0.78 | 0.434 | 0.501 |
| HRV change (last - first) | 0.46 | 0.502 | 0.891 | 0.39 | 0.536 | 0.894 | 0.01 | 0.931 | 0.971 | -0.54 | 2.88 | -0.19 | 0.851 | 0.851 |

**Table 5I.** Results from GLM tests of moderation. 'DV' refers to dependent variable of EDI from the post timepoint. 'Psych Moderator' and 'Physio Moderator' refer to either the psychological or physiological variable defined as the moderator. 'Tech x Stim x Moderator' refers to the model of a three-way interaction between the moderator and the group factors of Tech (MindGym vs VR) and Stimuli (Breathwork vs Rain). 'Tech x Moderator' refers to the model of a two-way interaction between the moderator and the group factor of Tech (MindGym vs VR). 'Stim x Moderator' refers to the model of a two-way interaction between the moderator and the group factor of Stimuli (Breathwork vs Rain). 'Moderator' refers to the effect of the moderator from the model of only main effects. 'F' refers to F-test from the GLM model. 'b' and 'SE' refer to the unstandardized beta coefficient and its standard error. 'Z' refers to the Z-test from the model. 'p.raw' and 'p.fdr' refer to the uncorrected p value and FDR-corrected p value.

#### Table 5J (BSE)

| DV | Tech x Stim x Moderator | | | Tech x Moderator | | | Stim x Moderator | | | Moderator | | | | |
|---|---|---|---|---|---|---|---|---|---|---|---|---|---|---|
| BSE | **F** | **p.raw** | **p.fdr** | **F** | **p.raw** | **p.fdr** | **F** | **p.raw** | **p.fdr** | **b** | **SE** | **Z** | **p.raw** | **p.fdr** |
| **Psych Moderator** | | | | | | | | | | | | | | |
| DPES | 0.81 | 0.371 | 0.888 | 0.57 | 0.452 | 0.974 | 2.60 | 0.110 | 0.671 | -0.15 | 0.18 | -0.83 | 0.409 | 0.767 |
| Immersion | 0.11 | 0.743 | 0.888 | 0.01 | 0.944 | 0.974 | 0.24 | 0.626 | 0.892 | -0.02 | 0.01 | -1.54 | 0.125 | 0.700 |
| IPQ (General) | 0.77 | 0.381 | 0.888 | 0.26 | 0.610 | 0.974 | 1.62 | 0.206 | 0.671 | -0.16 | 0.09 | -1.77 | 0.076 | 0.700 |
| IPQ (Involvement) | 0.86 | 0.355 | 0.888 | 5.47 | 0.021 | 0.315 | 2.30 | 0.132 | 0.671 | 0.03 | 0.04 | 0.84 | 0.404 | 0.767 |
| IPQ (Spatial Presence) | 0.09 | 0.761 | 0.888 | 0.69 | 0.409 | 0.974 | 0.40 | 0.528 | 0.888 | 0.01 | 0.05 | 0.16 | 0.876 | 0.898 |
| IPQ (Experienced Realism) | 1.26 | 0.263 | 0.888 | 0.80 | 0.374 | 0.974 | 2.07 | 0.153 | 0.671 | 0.04 | 0.04 | 0.97 | 0.330 | 0.767 |
| MODTAS | 0.01 | 0.912 | 0.965 | 1.32 | 0.253 | 0.974 | 0.96 | 0.329 | 0.705 | 0.00 | 0.01 | -0.13 | 0.898 | 0.898 |
| Openness | 0.09 | 0.770 | 0.888 | 0.02 | 0.901 | 0.974 | 0.06 | 0.815 | 0.892 | -0.01 | 0.04 | -0.31 | 0.760 | 0.898 |
| **Physio Moderator** | | | | | | | | | | | | | | |
| Motion (overall) | 0.09 | 0.766 | 0.888 | 0.56 | 0.457 | 0.974 | 0.03 | 0.862 | 0.892 | -0.01 | 0.02 | -0.29 | 0.769 | 0.898 |
| Breath Rate (first 2 min) | 0.18 | 0.677 | 0.888 | 0.03 | 0.862 | 0.974 | 0.12 | 0.726 | 0.892 | 1.00 | 79.00 | 1.26 | 0.265 | 0.767 |
| HR (first 1 min) | 0.00 | 0.965 | 0.965 | 1.80 | 0.184 | 0.974 | 1.25 | 0.268 | 0.671 | 0.00 | 0.02 | 0.15 | 0.884 | 0.898 |
| HRV (first 5 min) | 0.16 | 0.689 | 0.888 | 0.15 | 0.699 | 0.974 | 0.02 | 0.892 | 0.892 | 1.52 | 3.58 | 0.43 | 0.670 | 0.898 |
| Breath Rate change (last - first) | 0.52 | 0.473 | 0.888 | 0.02 | 0.895 | 0.974 | 1.37 | 0.246 | 0.671 | 0.06 | 0.04 | 1.48 | 0.140 | 0.700 |
| HR change (last - first) | 3.63 | 0.061 | 0.888 | 0.05 | 0.833 | 0.974 | 0.39 | 0.533 | 0.888 | -0.04 | 0.04 | -0.96 | 0.339 | 0.767 |
| HRV change (last - first) | 1.35 | 0.249 | 0.888 | 0.00 | 0.974 | 0.974 | 0.03 | 0.867 | 0.892 | -0.33 | 2.10 | -0.16 | 0.877 | 0.898 |

**Table 5J.** Results from GLM tests of moderation. 'DV' refers to dependent variable of BSE from the post timepoint. 'Psych Moderator' and 'Physio Moderator' refer to either the psychological or physiological variable defined as the moderator. 'Tech x Stim x Moderator' refers to the model of a three-way interaction between the moderator and the group factors of Tech (MindGym vs VR) and Stimuli (Breathwork vs Rain). 'Tech x Moderator' refers to the model of a two-way interaction between the moderator and the group factor of Tech (MindGym vs VR). 'Stim x Moderator' refers to the model of a two-way interaction between the moderator and the group factor of Stimuli (Breathwork vs Rain). 'Moderator' refers to the effect of the moderator from the model of only main effects. 'F' refers to F-test from the GLM model. 'b' and 'SE' refer to the unstandardized beta coefficient and its standard error. 'Z' refers to the Z-test from the model. 'p.raw' and 'p.fdr' refer to the uncorrected p value and FDR-corrected p value.
