## Supplementary material for "Contrasting cognitive, behavioral, and physiological responses to breathwork vs. naturalistic stimuli in reflective chamber and VR headset environments": S6 Tables A-C; Figure A.pdf

### S6: Follow-up Data

#### Table 6A (Descriptive Statistics)

| | | STAI | | POMS | | GA-VAS | |
|---|---|---|---|---|---|---|---|
| **Group** | **Timepoint** | **M** | **SD** | **M** | **SD** | **M** | **SD** |
| **MindGym (Breathwork)** | **Pre** | 34.96 | 8.43 | 9.59 | 23.33 | n/a | n/a |
| | **Post** | 30.18 | 7.74 | -0.86 | 18.51 | n/a | n/a |
| | **FollowUp** | 37.23 | 12.36 | 7.91 | 20.71 | 3.77 | 1.97 |
| **MindGym (Rain)** | **Pre** | 37.29 | 9.46 | 15.33 | 22.09 | n/a | n/a |
| | **Post** | 33.79 | 6.00 | 5.04 | 17.28 | n/a | n/a |
| | **FollowUp** | 39.63 | 11.61 | 10.54 | 17.39 | 4.25 | 2.19 |
| **VR (Breathwork)** | **Pre** | 34.10 | 7.44 | 7.05 | 19.11 | n/a | n/a |
| | **Post** | 29.10 | 4.67 | -1.90 | 13.15 | n/a | n/a |
| | **FollowUp** | 36.30 | 7.61 | 4.65 | 12.53 | 3.95 | 1.85 |
| **VR (Rain)** | **Pre** | 33.24 | 9.04 | 6.38 | 27.12 | n/a | n/a |
| | **Post** | 32.95 | 6.52 | 2.10 | 22.07 | n/a | n/a |
| | **FollowUp** | 36.24 | 8.23 | 8.62 | 15.68 | 3.67 | 1.77 |

**Table 6A.** STAI = State Trait Anxiety Scale; POMS = Profile of Mood States (Total Mood Disturbance); GA-VAS = Generalized Anxiety - Visual Analog Scale; M = Mean SD = standard deviation

#### Figure 6A (Descriptive statistics by Timepoints)

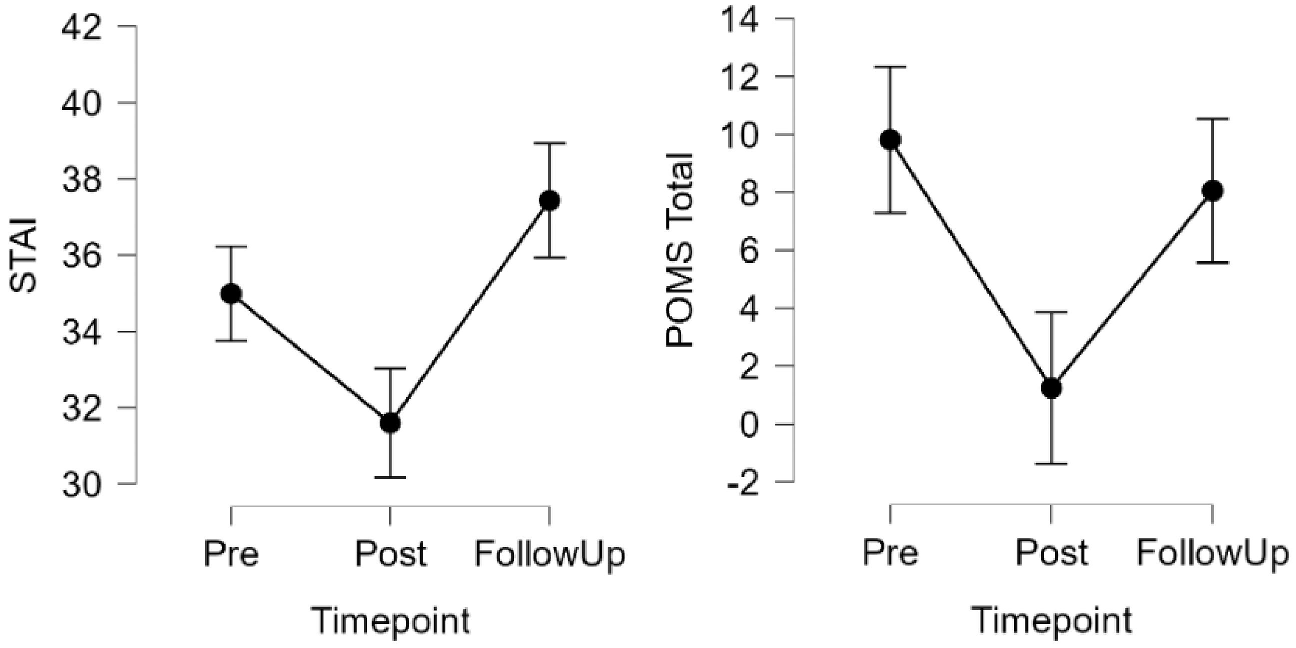

#### Table 6B (Repeated measures ANOVA)

| | Timepoint | | | | | | Timepoint x Tech | | | | | |
|---|---|---|---|---|---|---|---|---|---|---|---|---|
| **DV** | **df1** | **df2** | **F** | **p.raw** | **p.fdr** | **$\eta^2_G$** | **df1** | **df2** | **F** | **p.raw** | **p.fdr** | **$\eta^2_G$** |
| **STAI** | 2.00 | 172.00 | 17.48 | < .001 | 0.004 | 0.073 | 2.00 | 170.00 | 0.33 | 0.722 | 0.772 | 0.001 |
| **POMS** | 2.00 | 172.00 | 12.55 | < .001 | 0.004 | 0.035 | 2.00 | 170.00 | 0.65 | 0.525 | 0.759 | 0.002 |
| | **Timepoint x Stim** | | | | | | **Timepoint x Tech x Stim** | | | | | |
| **DV** | **df1** | **df2** | **F** | **p.raw** | **p.fdr** | **$\eta^2_G$** | **df1** | **df2** | **F** | **p.raw** | **p.fdr** | **$\eta^2_G$** |
| **STAI** | 2.00 | 170.00 | 1.24 | 0.293 | 0.586 | 0.006 | 2.00 | 166.00 | 0.42 | 0.664 | 0.772 | 0.002 |
| **POMS** | 2.00 | 170.00 | 0.21 | 0.811 | 0.811 | 0.001 | 2.00 | 166.00 | 0.57 | 0.569 | 0.759 | 0.002 |

**Table 6B** DV = dependent variable; STAI = State Trait Anxiety Scale; POMS = Profile of Mood States (Total Mood Disturbance); Timepoint = main effect model (within-subjects: pre, post, follow up); Timepoint x Tech = interaction model (between-groups: MindGym vs VR); Timepoint x Stim = interaction model (between-groups: Breathwork vs Rain); Timepoint x Tech x Stim = three-way interaction model; F = F-test with df1, df2 (degrees of freedom); P.RAW/p.fdr = uncorrected/FDR-corrected p-values

#### Table 6C (Post hoc comparisons)

| | Post hoc tests by Timepoint (STAI) | | | | | |
|---|---|---|---|---|---|---|
| **Timepoints** | **ΔM** | **SE** | **df** | **t** | **Cohen's d** | **p.fdr** |
| **Post - Pre** | -3.39 | 0.99 | 172.00 | -3.42 | -0.40 | 0.002 |
| **FollowUp - Pre** | 2.45 | 0.99 | 172.00 | 2.47 | 0.29 | 0.015 |
| **FollowUp - Post** | 5.84 | 0.99 | 172.00 | 5.89 | 0.68 | < .001 |
| | **Post hoc tests by Timepoint (POMS)** | | | | | |
| **Timepoints** | **ΔM** | **SE** | **df** | **t** | **Cohen's d** | **p.fdr** |
| **Post - Pre** | -8.58 | 1.81 | 172.00 | -4.74 | -0.44 | < .001 |
| **FollowUp - Pre** | -1.76 | 1.81 | 172.00 | -0.97 | -0.09 | 0.332 |
| **FollowUp - Post** | 6.82 | 1.81 | 172.00 | 3.77 | 0.35 | < .001 |

**Table 6C** STAI = State Trait Anxiety Scale; POMS = Profile of Mood States (Total Mood Disturbance); ΔM/SE = mean difference/standard error between timepoints; t, df = t-test statistic and degrees of freedom; Cohen's d = effect size; p.fdr = FDR-corrected p-value
